# Human spinal interneurons repair the injured spinal cord through synaptic integration

**DOI:** 10.1101/2024.01.11.575264

**Authors:** Lyandysha V. Zholudeva, Tara Fortino, Ayushi Agrawal, Olaia F. Vila, Maggie Williams, Todd McDevitt, Michael A. Lane, Deepak Srivastava

## Abstract

Advances in cell therapy offer promise for some of the most devastating neural injuries, including spinal cord injury (SCI). Endogenous VSX2-expressing spinal V2a interneurons have been implicated as a key component in plasticity and therapeutically driven recovery post-SCI. While transplantation of generic V2a neurons may have therapeutic value, generation of human spinal V2a neurons with rostro-caudal specificity and assessment of their functional synaptic integration with the injured spinal cord has been elusive. Here, we efficiently differentiated optogenetically engineered cervical V2a spinal interneurons (SpINs) from human induced pluripotent stem cells and tested their capacity to form functional synapses with injured diaphragm motor networks in a clinically-relevant sub-acute model of cervical contusion injury. Neuroanatomical tracing and immunohistochemistry demonstrated transplant integration and synaptic connectivity with injured host tissue. Optogenetic activation of transplanted human V2a SpINs revealed functional synaptic connectivity to injured host circuits, culminating in improved diaphragm activity assessed by electromyography. Furthermore, optogenetic activation of host supraspinal pathways revealed functional innervation of transplanted cells by host neurons, which also led to enhanced diaphragm contraction indicative of a functional neuronal relay. Single cell analyses pre- and post-transplantation suggested the *in vivo* environment resulted in maturation of cervical SpINs that mediate the formation of neuronal relays, as well as differentiation of glial progenitors involved in repair of the damaged spinal cord. This study rigorously demonstrates feasibility of generating human cervical spinal V2a interneurons that develop functional host-transplant and transplant-host connectivity resulting in improved muscle activity post-SCI.

## [INTRODUCTION]

Spinal cord injury (SCI) remains a devastating and often unpredictable event with no curative therapeutic interventions. While current treatments for SCI (e.g., rehabilitation, neuromodulation) can improve patient quality of life^1–5^, functional improvements via these approaches are reliant on harnessing plasticity in neural tissue spared by injury, and leave the underlying tissue damage untreated. In contrast, rapid advances in regenerative medicine are providing promising pharmacological, gene, and cell-based strategies that enhance neural repair^6^. However, such approaches have so far achieved only partial functional improvement, perhaps because they too often focus on regeneration or restoration of lost pathways, rather than building on intrinsic neuroplasticity. Efforts to combine regenerative medicine with intrinsic plasticity may offer a more successful approach to SCI therapy.

An important development has been the realization that spinal interneurons (SpINs) may be the gateway to plasticity and recovery after injury and disease in the spinal cord^7,8^. SpINs represent the most dominant population of neurons in the spinal cord, modulating input and output within and between networks, and are not only essential to normal functional spinal networks, but also key components of plasticity after injury or disease^7^. While there are several cardinal classes of interneurons^7,9^, and likely hundreds of interneuronal phenotypes overall, VSX2 (a.k.a. CHX10)-expressing V2a interneurons have been identified as a potential candidate for spontaneous^10^ and therapeutically mediated^1^ plasticity and recovery. Regional identity of transplanted cells may be important for targeted restoration of specific circuits^11–13^, including rostro-caudal differentiation of SpINs based on the level of SCI. However, transplantation of well-defined cervical human SpINs (e.g., V2a interneurons), and rigorous evaluation for functional integration has not been performed.

Here, we differentiated a population of the VSX2-positive V2a SpINs with cervical identity from optogenetically engineered human induced pluripotent stem cells (hiPSCs), and demonstrated their functional integration with the injured spinal cord. Specifically, we harnessed the relative simplicity of a cervical spinal respiratory network—the phrenic circuit—to demonstrate the ability of V2a SpINs to form functional, host-donor and donor-host synaptic connectivity. Leveraging advanced optogenetics and multi-unit electrophysiology, we show that transplanted human SpINs can provide and receive functional synaptic connections with injured motor networks resulting in muscle contraction. These experiments provide the first in-depth anatomical and electrophysiological assessment of functional synaptic integration between the injured host spinal cord and donor human SpINs, and vice-versa.

## [RESULTS]

### V2a spinal interneurons can be engineered from human stem cells

We engineered human V2a spinal interneurons (SpINs) from an optogenetic channelrhodopsin-2-expressing human induced pluripotent stem cell (hiPSC) line^14^ using a combination of small molecules that neuralized, ventralized, and caudalized the cells over time (Fig 1a-b). While prior studies have established protocols to drive the differentiation of V2a neurons^15^, we modified these methods to caudalize the cells toward a spinal fate. To achieve this, we added the Wnt agonist (CHIR) and tested several concentrations to achieve rapid expression of cervical spinal cord transcription factors (HOX genes, Fig 1c) and neuronal genes, but limit expression of glial or retinal genes (Fig 1d). This was best achieved with 4 uM CHIR. With this optimized protocol, we generated a large batch of light-activatable human SpINs enriched for the V2a population by replating (without cryopreservation) at day 15 of differentiation (Fig 1e-h), and evaluated their electrical activity *in vitro* using multielectrode arrays (MEA, Fig 1i-o), recording spontaneous and activated (blue light) electrical activity over time (see Fig 1l raster plot, movies available in Supplementary Information). Spontaneous activity (number of spikes, Fig 1m) as well as neuronal synchronization (number of bursts, Fig 1n) increased with time *in vitro*, indicating neuronal maturation and network formation^16^. Cells responded to blue light stimulation in synchrony with the frequency used (Fig 1o). Blue light stimulation of the cells while plated on MEAs containing spinal motor neurons demonstrated the capacity for these SpINs to form functional connections with motor neurons *in vitro* (Extended Data Fig 1a-d).

**Figure 1.**
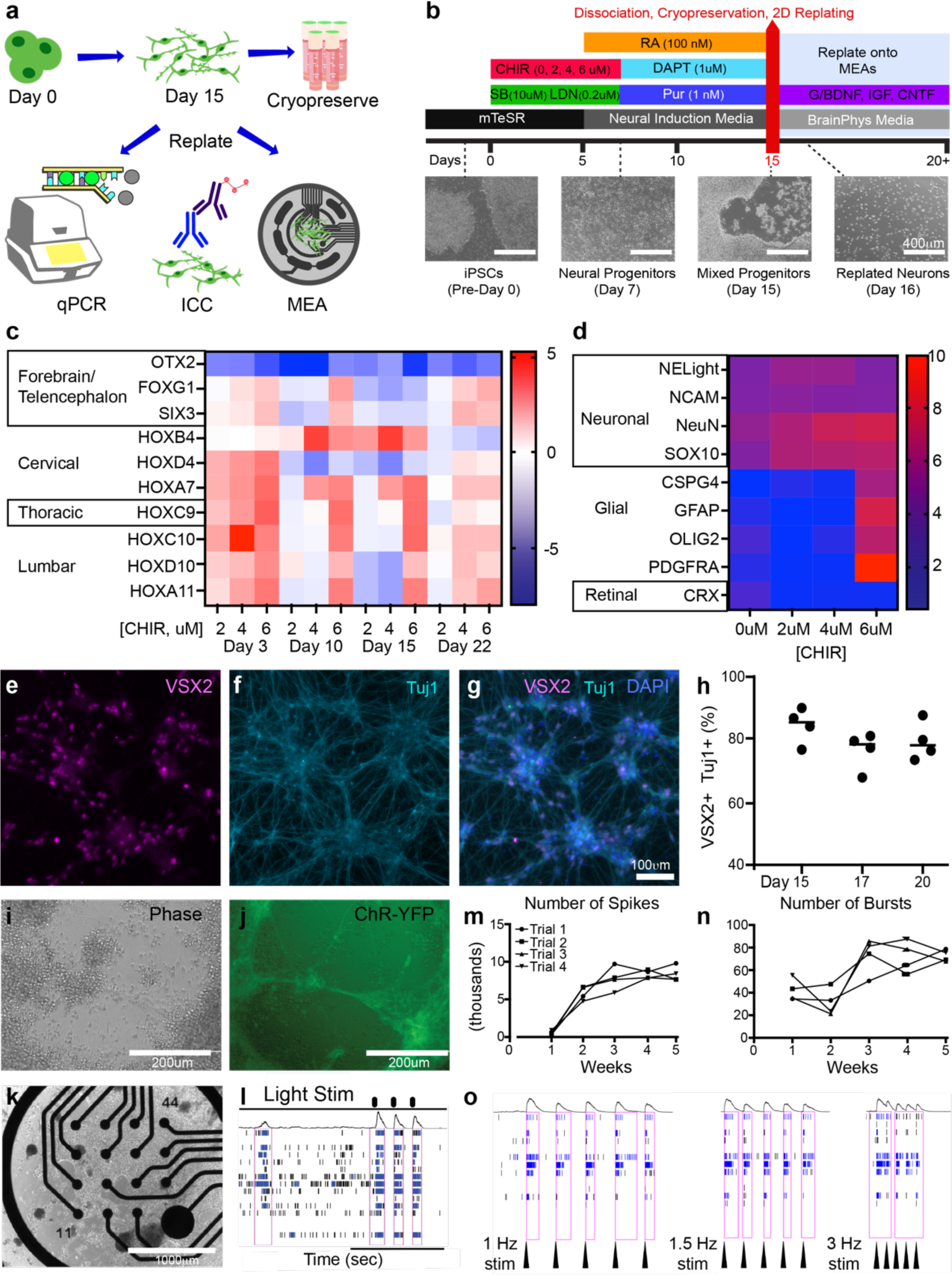
Human V2a spinal interneurons can be differentiated from iPSCs. A 15 day protocol was used to differentiate and characterize human spinal interneurons (SpINs), from human induced pluripotent stem cells (hiPSCs; a-b). Analysis of rostro-caudal gene expression using qPCR in cells treated with 2, 4, or 6 uM of CHIR, normalized to 0 uM CHIR (c), showed that 4uM resulted in the greatest expression of cervical genes (e.g., HOXB4) at Day 15, while maintaining neuronal gene expression (d, normalized to hiPSCs). Immunocytochemistry of cells after replating on Days 15, 17, and 20 revealed that around 80% of cells co-labeled for VSX2 (e,g,h) and the neuronal marker Tuj1 (f,g,h). Optogenetic hiPSC line with YFP reporter (ChR-YFP; i,j) was used to generate and replate V2a SpINs onto multielectrode arrays (MEAs, k) to record their electrical activity (l), demonstrating functional control of the cells *in vitro*. The number of spontaneously recorded spikes (m) and network bursting (n) increased over time, consistently across biological replicates (trials) over 5 weeks *in vitro*. Light stimulation of cells (1, 1.5 and 3 Hz) while recording from the electrodes resulted in significant enhancement of detectable activity (o). Arrowheads in (o) point to stimulation events.

To characterize the transcriptomic profile of the day 15 hiPSC-derived SpINs, we performed single-cell RNA-sequencing (scRNA-seq) of freshly thawed, “off the shelf” samples, from the same batch to be used for transplantation. We sequenced and analyzed 15,361 cells, which resulted in five distinct clusters (designated 0-4) after quality control filtering (Fig 2a), with majority of the cells expressing either progenitor V2a marker VSX1 or the post-mitotic marker for V2a, VSX2 (Fig 2b). The top 10 differentially expressed genes for each cluster were used to distinguish the transcriptional profiles of the five clusters (Fig 2c, Extended Data Table 1) and individual inspection of the gene lists and databases^17,18^ were used to assign identities to each cluster (Fig 2a, c, with additional feature plots shown in Extended Data Fig 2). We also evaluated expression of homeobox (HOX) genes characteristic of anterior (forebrain) versus posterior (spinal cord) neurons, which revealed the transplantable cells were primarily of cervical spinal cord phenotype (Fig 2d). Genes typically expressed in the forebrain (FOXG1, LMX1, FOXA2, etc.) were absent, whereas HOX genes expressed in the cervical and thoracic spinal cord were expressed in all clusters (Fig 2d, Extended Data Fig 3).

**Figure 2.**
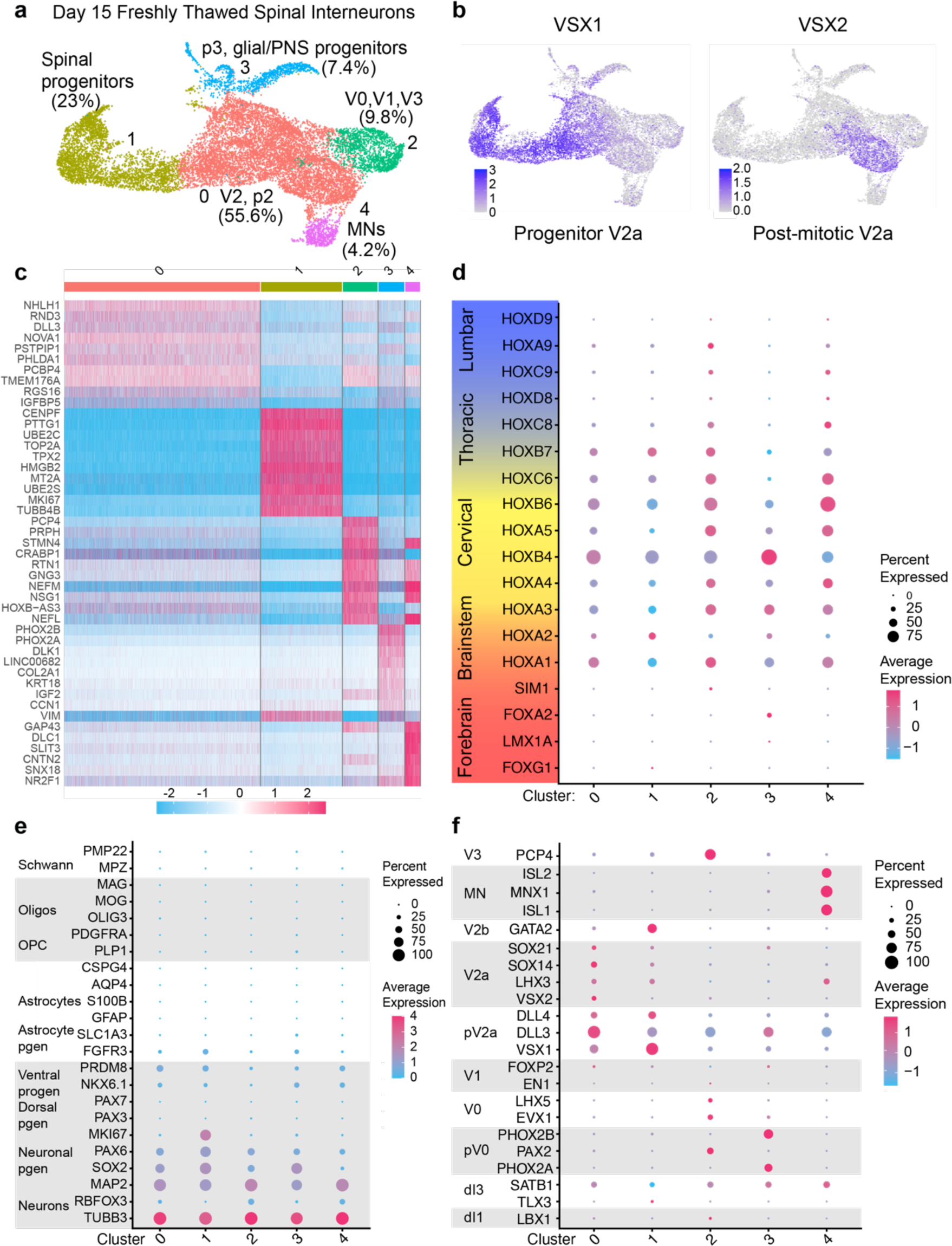
Single cell RNA sequencing of human V2a SpINs. (a) A UMAP plot of scRNAseq data from Day 15 human iPSC-derived cells thawed from cryopreservation revealed that the majority of cells were either V2a SpINs (55.6%, cluster 0) or progenitor cells (30.4%, cluster 1 & 3). A small number of other ventrally derived spinal neurons were also detected: 9.8% V0/V1/V3 SpINs (cluster 2), and 4.2% motor neurons (MNs; cluster 4). (b) UMAPs showing relative distribution of VSX1-expressing progenitor V2as (left) and VSX2-expressing post-mitotic V2as (right). (c) Heatmap plot of top expressing genes in each of the clusters. (d) Analysis of HOX gene expression revealed the relative rostrocaudal identity of the cells to be largely cervical. (e) Dot plot of neuronal and glial genes revealed that most cells were neuronal or neural progenitors. (f) Dot plot of developmental transcription factors enriched in spinal cardinal classes.

Normalized average expression of neuronal, glial, and developmental transcription factor-encoding genes revealed that all clusters were comprised of spinal neurons or their progenitors (Fig 2e), and most expressed genes associated with V2a SpINs or motor neurons (MNs; Fig 2f). These data showed that the vast majority of the cells were neuronal (TUBB3+, RBFOX3), and predominantly progenitors (SOX2, PAX6), with the majority being ventrally derived (PRDM8 and NKX6.1) (Fig 2e). Dorsal progenitors expressing either PAX3 or PAX7^17^ comprised <1% of the population (Extended Data Table 2). Less than 1% of the cells expressed any of the genes commonly used to identify glial cells, such as astrocytes (e.g., AQP4, S100B, GFAP), oligodendrocytes (OLIG3, MOG) or Schwann cells (PMP22, MPZ; Figure 2e, Extended Data Table 2). We also assessed expression of canonical transcription factors (TFs) used to identify classes of SpINs, revealing that nearly 60% of cells expressed VSX1, indicative of the V2a progenitor population, and 40.3% of cells were PRDM8+, marking V2 neurons and their progenitors. A significant number of cells also expressed more mature markers of V2a cells (SOX14, LHX3 and VSX2, Extended Data Table 2). Importantly, <1% of cells expressed Engrailed-1, a marker of the inhibitory V1 population (Extended Data Table 2). Taken together, this characterization indicated that our engineered hiPSCs comprise cervical spinal cord neurons and their progenitors, and are predominantly of V2a SpIN identity.

### Transplanted V2a SpINs functionally integrate with injured phrenic circuit

A lesion cavity typically forms in the damaged spinal cord within weeks (sub-acutely) after traumatic SCI in humans and in pre-clinical animal models. This creates an enclosed intraspinal environment into which transplanted cells can be safely delivered. To test the ability of transplanted human V2a SpINs to functionally integrate with the injured host spinal cord, we used an adult Sprague Dawley rat model of cervical SCI (C3/4 contusion injury) and transplanted Day 15 optogenetically engineered hiPSC-derived SpINs directly into the injury site sub-acutely (one week) after injury (Fig 3a-b). We evaluated three different doses (250k, 500k, 1 million cells/animal; Extended Data Fig 4a-d), and histological assessment revealed that survival and confluency at 1 month were best achieved with 1 million cells/animal (125k cells per microliter). Subsequent studies described here are based on this dose, using cryopreserved cells that were thawed and transplanted on the same day, which provided methodological consistency and clinical relevance for ultimate “off the shelf” applications.

**Figure 3.**
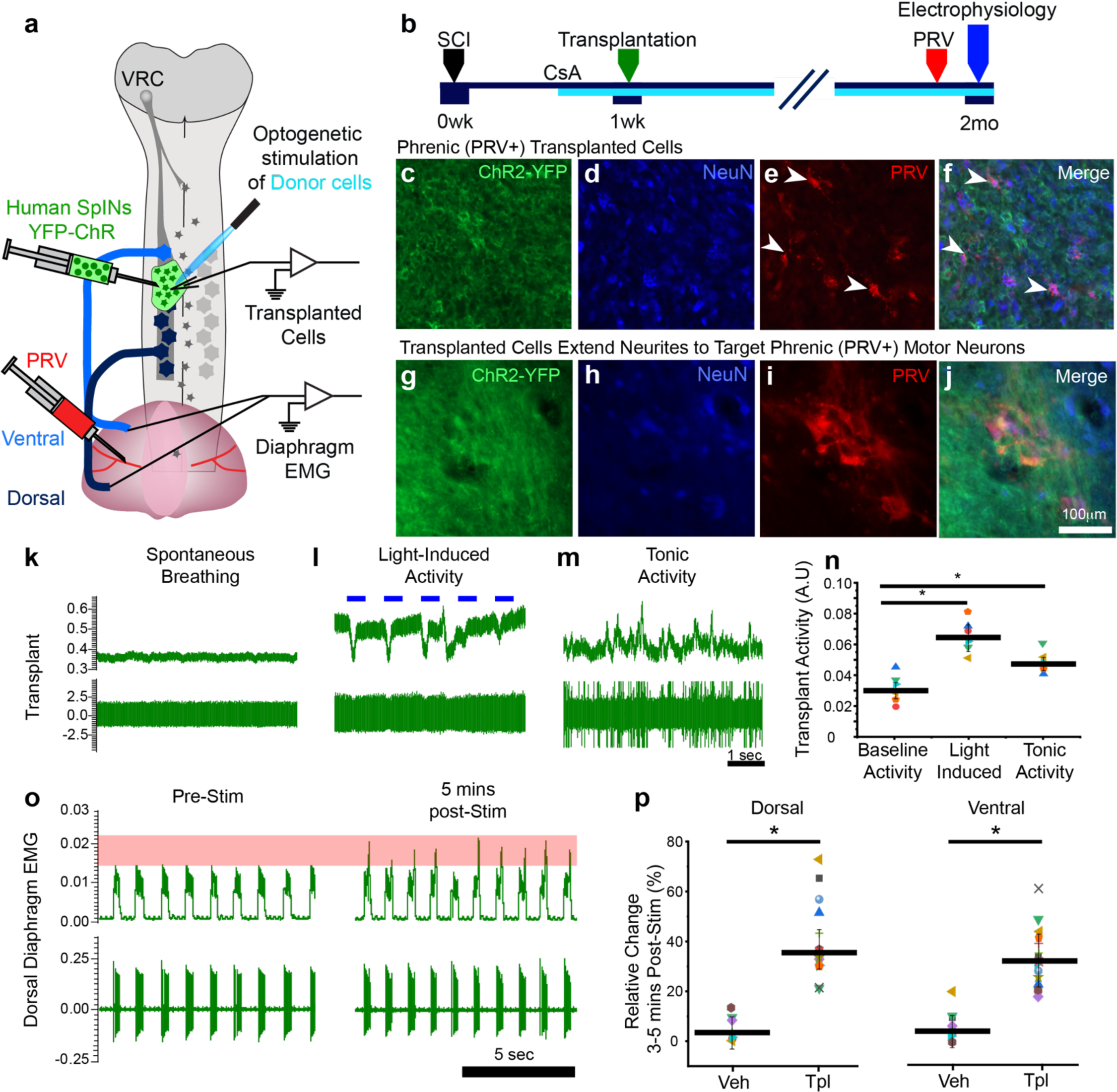
Transplanted V2a SpINs functionally integrate with injured host motor network. Schematic diagram of experimental design (a) and timeline (b). Transneuronal tracing with pseudorabies virus (PRV) demonstrates anatomical connectivity of transplanted (ChR2-YPF+, c) neurons (NeuN+, d) with host phrenic circuit (PRV+, e and merge in f) with transplanted neurites extending to PRV+ phrenic motor pool (g-j). Light stimulation of donor neurons revealed immediate bursting within donor neurons (green traces k-m, quantification in n). Example of diaphragm electromyography trace prior to stimulation and 5 minutes post-stimulation (o), with percent response to stimulation quantified over 40 seconds (p) from dorsal site of the diaphragm (left) and ventral site of the diaphragm (right). Veh: vehicle control recipient, Tpl: transplant recipient. ANOVA, Tukey’s post-hoc, *p<0.05.

Two months after transplantation, histology from transplant recipients revealed that the donor human cells survived, filled the injury site, and anatomically integrated with the host injured cord (Extended Data Fig 4e-h). Donor cells were confluent with injured host tissue, with robust vascularization and serotonergic innervation (Extended Data Fig 4f-h).

To determine whether donor neurons synaptically integrated with denervated spinal networks caudal to the injury, we delivered a transsynaptic neuroanatomical tracer—pseudorabies virus (PRV)—to the diaphragm. We observed PRV-positive transplanted human SpINs 72 hours post-infection, indicating synaptic integration with the injured phrenic circuit (donor-to-host connectivity; Fig 3c-f), with neurites extending to target phrenic motor neurons within the spinal cord (Fig 3g-j). PRV tracing revealed robust donor-to-host synaptic connectivity in the majority of animals (83%), with few instances of animals with limited connectivity (Extended Data Fig 5).

To test the function of these donor-to-host connections, the spinal cord at the site of the transplant was re-exposed 2 months post-transplantation and donor human V2a SpINs were stimulated with blue light (Extended Data Fig 6a-c). A multi-unit electrode (Carbostar) was placed into the transplant, while tungsten electrodes were implanted into the ventral and dorsal ipsilateral-to-injury hemidiaphragm. Multi-unit donor cell and diaphragm activity were recorded during spontaneous breathing (Fig 3k) for at least 10 minutes prior to blue light stimulation (3 minutes, 1Hz, see Methods). Stimulation induced immediate increase in transplant activity (Fig 3l), and this activity persisted after stimulation was stopped (Fig 3m,n). A robust increase in diaphragm activity was seen 3-5 minutes after the 3-minute stimulation protocol was ceased, as measured from both dorsal and ventral hemidiaphragm sites (Fig 3o, p). This effect could be repeated after periods of rest (Extended Data Fig 7).

### Supraspinal host axons functionally integrate with transplanted V2a SpINs

As expected, we observed robust innervation of donor tissue from serotonergic host axons (5HT, Extended Data Fig 4e-h), which modulate respiratory networks^19,20^. We also observed bulbospinal axons from the ventral respiratory column (VRC), which have a lower propensity for spontaneous growth, innervating donor cells two months post-transplantation. To test function of VRC input, we injected AAV1 containing the Syn-Chrimson-tdTomato^21^ gene cassette into brainstem reticular nuclei one week prior to the terminal experiment (Fig 4a,b and Extended Data Fig 8a-c). Chrimson (red-light activated channelrhodopsin) and tdTomato positive bulbospinal axons extending into the transplant could then be functionally and anatomically assessed, respectively. TdTomato positive axons were observed throughout the transplant one-week post-injection (Fig 4c-d). TdTomato positive puncta were closely juxtaposed to synaptophysin proteins and phrenic (PRV+) transplanted cells (Fig 4c-h). Multi-unit activity from transplanted cells was recorded prior to (Fig 4i), during (Fig 4j) and post red-light stimulation (Fig 4k). Transplant activity synchronized to host-stimulation in the presence of light (1Hz stimulation, Fig 4j), and in the animals that responded to initial light stimulation this effect could be repeated throughout the duration of the recording, providing evidence of host-to-donor functional synapses. Transplant activity persisted for 1-5 minutes after light-stimulation ceased (Fig 4l). These results demonstrate that reticular neurons extended axons into the region of transplanted human SpINs and formed functionally viable connections.

**Figure 4.**
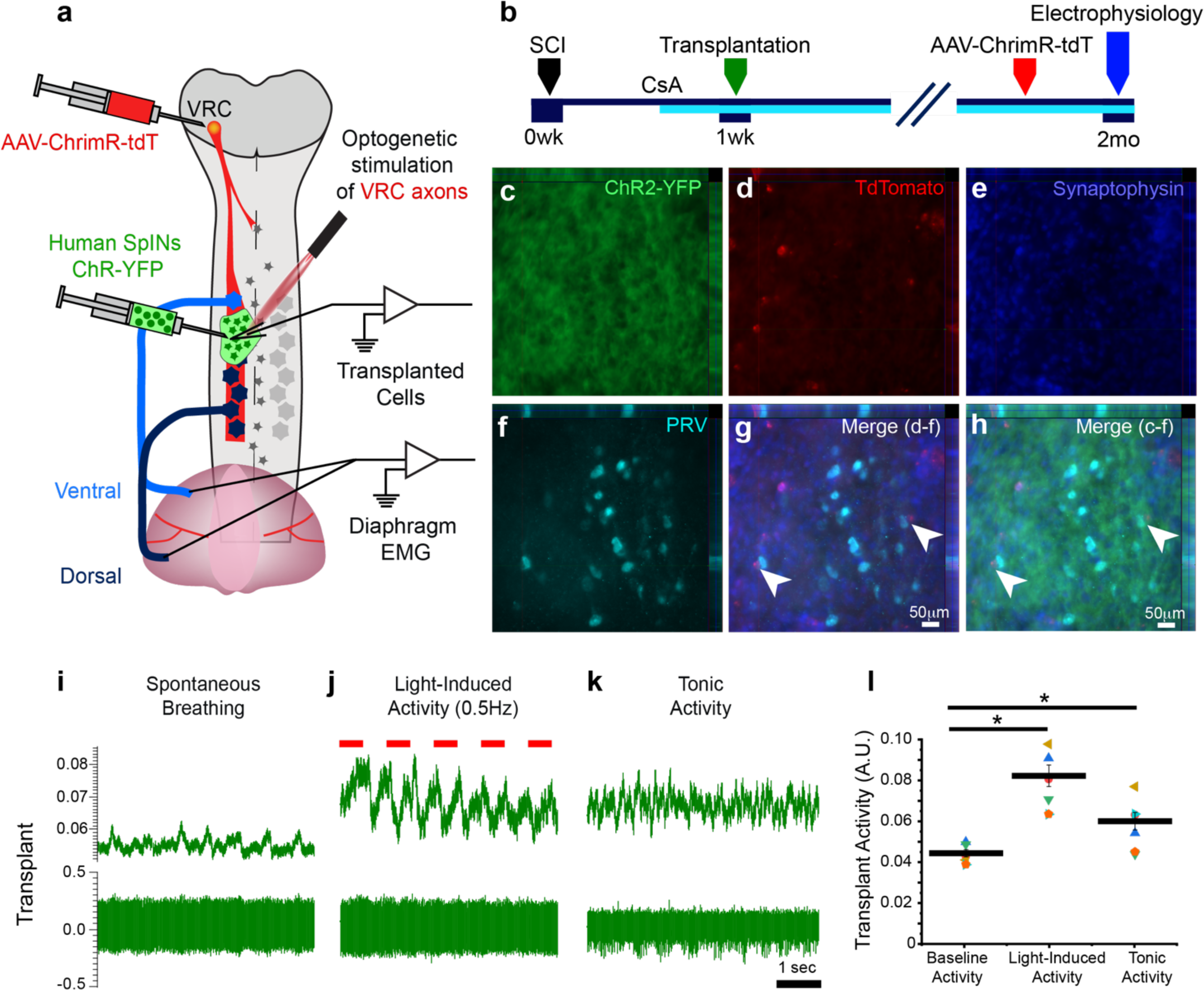
Injured host axons functionally innervate transplanted human V2a SpINs. Schematic diagram of experimental design (a) and timeline (b). Coronal section through transplant epicenter shows YFP+ transplanted cells (c) with abundant expression of ChrimsonR-tdTomato+ punctae throughout the transplant (d), forming synapses (synaptophysin+, e) with PRV-positive transplanted cells (f). (g) Merge of d-f; (h) merge of c-f. Sample traces of terminal electrophysiological recordings from the transplant during spontaneous breathing (i), during active stimulation (j) and post-stimulation (k) with red light. Quantification of activity during baseline, during, and post-redlight stimulation over 40 seconds (l). ANOVA, Tukey’s post-hoc, *p<0.05.

### Transplanted human V2a SpINs improve respiratory recovery

Two months following transplantation, animals underwent diaphragm electromyography (dEMG) while spontaneously breathing room air (eupneic breathing, considered ‘baseline’ condition) and during respiratory challenge (exposed to 10% oxygen (hypoxia) or 7% carbon dioxide (hypercapnia) inhaled gas) (Fig. 5a-c). Respiratory challenge is commonly used to assess an animal’s ability to respond to increased respiratory demand. Electrodes were surgically implanted into dorsal and ventral sites of the diaphragm to measure muscle activity under baseline and challenge conditions (Fig 5c for example traces). Given the documented somatotopic innervation of the diaphragm^22,23^, the ventral and dorsal recording sites are associated with innervation from the rostral and caudal phrenic motor pool, respectively. Following a C3/4 contusion injury, a dorsal muscle recording (innervated by phrenic motor neurons below C4-level) corresponds to the network that has been denervated below the injury, and muscle function can be assessed with maximum amplitude of diaphragm output (Fig 5c).

**Figure 5.**
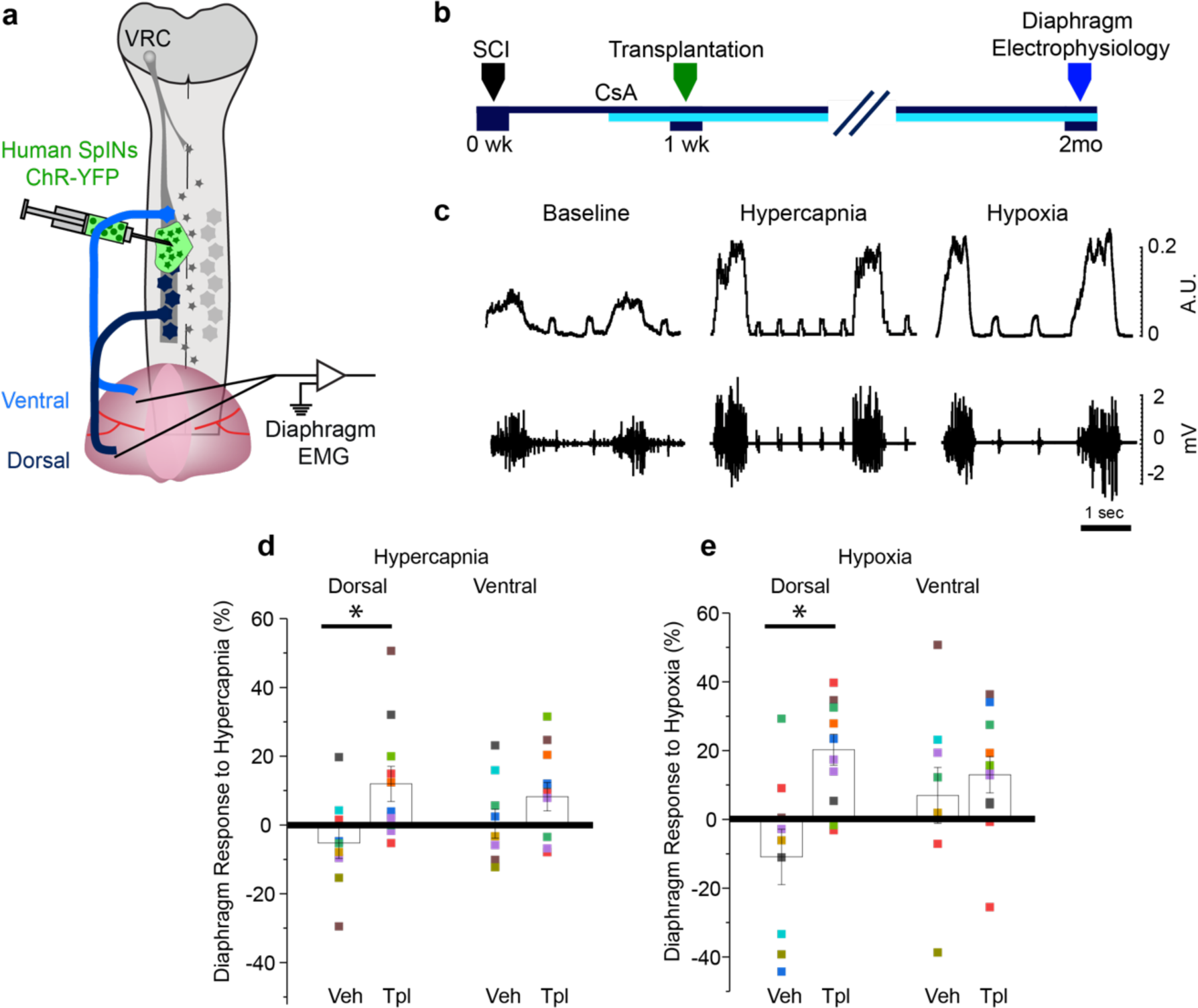
Human V2a SpINs promote respiratory recovery. Schematic diagram of experimental design (a) and timeline (b) for assessing the contribution of transplanted human V2a SpIN to respiratory recovery as measured with diaphragm muscle electromyography (EMG). Diaphragm EMG were measured from animals breathing room air (baseline) and under challenged conditions (hypercapnia or hypoxia) as shown in the raw recording (bottom) and integrated traces (c). Average percent response of dorsal (left) and ventral (right) diaphragm activity ipsilateral to the injury, to either hypercapnic (d) or hypoxic (e) challenge (increase in activity relative to baseline) is represented for vehicle and transplant-recipients. ANOVA Tukey’s post-hoc, *p<0.05).

Diaphragm activity was statistically similar between vehicle controls and transplant recipients during baseline breathing. However, when challenged with hypercapnia or hypoxia, activity of dorsal diaphragm was greater in transplant recipients. In fact, the percent response to respiratory challenge was significantly greater in transplant recipients compared to that in the vehicle-treated control animals (ANOVA, Tukey post-hoc; p<0.05; Fig 5d-e). The extent to which output increased with challenge was also greatest in the dorsal region of the diaphragm, innervated by motor neurons caudal to the transplant. This result supports the fact that donor human V2a neurons integrate with the damaged phrenic network caudal to injury. Notably, most vehicle-treated animals showed a paradoxical decrease in diaphragmatic contraction in response to either challenge. This reduced muscle activity during challenge is consistent with muscle fatigue and a life-threatening form of respiratory insufficiency after injury, which can lead to respiratory arrest. While the relative increase in muscle activity seen in transplant recipients was quantitatively modest, the fact that fewer animals in this treated group showed a negative response to challenge (only 18% to hypoxia, 27% to hypercapnia), reflects a biologically significant and impactful improvement in outcome.

### Unbiased characterization of transplanted human V2a SpINs reveals donor cell plasticity post-transplantation

Given the evidence that transplanted human V2a SpINs functionally integrated with injured rodent networks, we used single-nuclear RNA-sequencing (snRNA-seq) to determine how transplanted cells differentiated and matured after transplantation into a SCI. Analysis of 8,765 nuclei isolated from the area of injury 2 months post-transplantation resulted in twenty distinct clusters (designated 0-19) (Fig 6a) after quality control filtering. The differentially expressed genes of each cluster (Extended Data Table 3) and individual inspection of genes were used to assign cell-type identities to each of the clusters (Fig 6a-b, with additional feature plots shown in Extended Data Fig 9). snRNA-seq data were aligned to the human reference genome and screened for Y-chromosome genes (e.g., USP9Y, Fig 6b), as the donor hiPSC line was derived from a male, while the rat recipients were females. Approximately half of the transplanted cells expressed neuronal genes (e.g., RBFOX3) and the remainder expressed glia-related genes (e.g., GFAP) (Fig 6c, e). Among the neuronal cells derived from the SpINs, we observed excitatory (e.g. SLC17A6 (VGLUT2)), inhibitory (e.g., PAX2) and neuromodulatory (e.g., non-peptidergic and peptidergic) neurons (Fig. 6b,d and Extended Data Fig. 10). Within these, both dorsal and ventral spinal neuronal markers were present (Extended Data Fig. 11; Extended Data Table 4). The excitatory spinal interneurons (VGLUT2-positive) represent the expected derivatives of VSX2-positive transplanted cells, and HOX gene expression reflected a cervical identity, further supporting maturation of cervical V2a neurons (downregulation of VSX2 in V2a neurons in this region^24^). The emergence of inhibitory interneurons was surprising, and suggests inherent neuroplasticity of donor cells and that the injured spinal cord environment may influence this plasticity.

**Figure 6.**
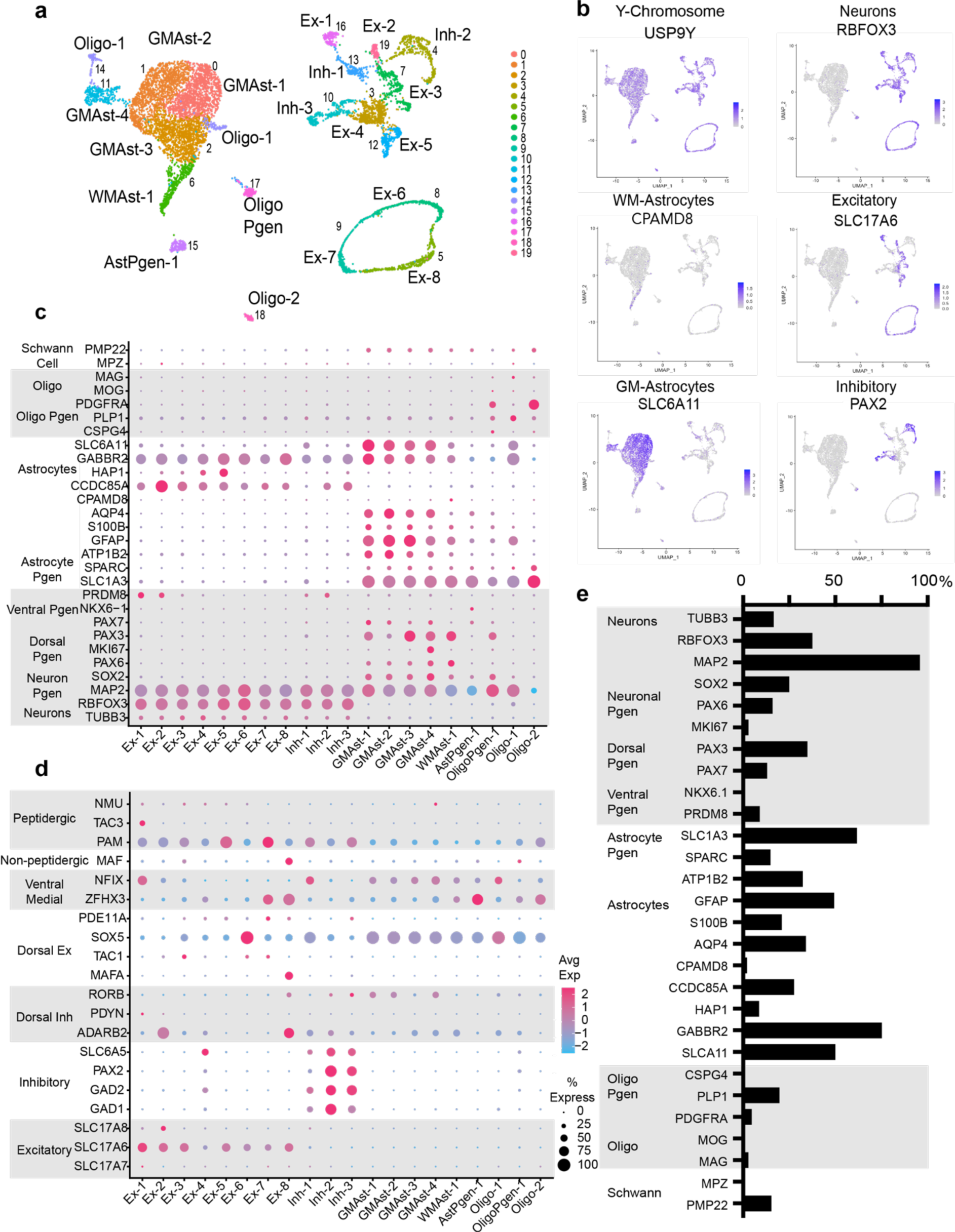
Single nuclear RNA sequencing of human V2a SpINs post-transplantation. A UMAP plot of RNA sequenced nuclei from animals that received SCI and transplants (n=3, 2 months post-transplant) shows 20 emergent clusters with approximately 50/50 split between glial and neuronal cells (a). UMAPs showing expression of Y-Chromosome-specific genes to ensure transplant versus host identity, as well as general markers for astrocytes and neurons (b). Dot plot of commonly-expressing genes in neuronal and glial cells (c) and neuronal subtypes (d) shows diversity of phenotypes 2-months post-transplantation. Relative percent expression of general markers for neuronal and glial phenotypes (e).

Despite the absence of astrocytes in the donor cells at the time of transplantation, approximately half of the human cells retrieved in the transplanted area expressed astrocytic markers (Fig. 6b and Extended Data Fig. 12). This suggests that some of the progenitors transplanted differentiate into astrocytes in the injured environment, though few oligodendrocytes were observed. Of donor cells that became astrocytes, the majority expressed genes associated with grey matter astrocytes (Extended Data Fig. 12). Furthermore, despite 10% of cells introduced being motor neurons, less than 1% of the cells 2 months later had motor neuron gene expression (Extended Data Fig 13a). Overall, the transplanted SpINs appeared to undergo further maturation *in vivo* and retained rostral identity (Extended Data Fig 13b), consistent with the integration with host tissues and functional improvement observed.

## [DISCUSSION]

Here we demonstrate that human V2a cervical SpINs can be engineered from hiPSCs, successfully transplanted into the injured spinal cord, and functionally integrate with damaged networks. The transplanted cells can both accept functional synaptic connections from the host (e.g., brainstem-to-transplant connectivity) as well as form functional synaptic connections with damaged and denervated host networks (transplant-to-phrenic network connectivity). By directing the differentiation of cervical V2a SpINs from an engineered optogenetic hiPSC line, we were able to selectively activate the transplanted cells post-transplantation while recording from the host phrenic circuit, assessing the transplants’ functional contribution to muscle recovery post-SCI. This quantitative evaluation of muscle activity provides a more rigorous and direct assessment of contribution to outcomes than the indirect behavioral measures that are typically used to assess motor recovery. Similarly, optogenetically activating host input to donor V2a SpINs revealed functional host-to-transplant connectivity. This rigorous evaluation of a functional host-donor-host relay using human cells in a model of clinically relevant, contusion cervical SCI definitively demonstrates the impact and anatomical mechanism by which transplanted cells can contribute to neural repair and recovery.

The transplantation of neural stem/progenitor cells for neural repair has offered new hope for therapeutic repair of the injured spinal cord^6,25^, providing the neuronal and glial building blocks needed for tissue repair. However, despite pre-clinical success and clinical translation, limitations persist. A major issue faced by prior work has been that despite our rapidly growing knowledge of neuronal cell phenotypes, and the identification of SpINs involved in functional recovery, existing transplant approaches use an undefined population of neuronal and glial progenitors, often with some stem cell components. To address this, we have developed a cell product enriched for therapeutic V2a SpINs with specified rostral-caudal identity, and leveraged single-cell approaches to transcriptionally define the hiPSC-derived cells pre- and post-transplantation.

The present data show that the transplanted neurons are not only innervated by the host, and functionally integrated with the host phrenic network, but also contribute to measurable improvements in diaphragm function after SCI. This improved function is seen most under conditions of increased respiratory drive, which might increase activation of donor neurons similar to optogenetic stimulation. This is a valuable, clinically relevant improvement, which may provide increased respiratory function at times of increased demand (e.g., during exertion, at altitude, during respiratory illness, etc.), and when individuals are potentially otherwise at risk of respiratory arrest. It may also provide an opportunity to further entrain muscle activity and strengthen the underlying neural networks that control them with additional therapeutic approaches (e.g., neural stimulation or activity-based therapies). Notably, there is some variability observed in the extent of connectivity between donor neurons and the host phrenic network, which suggests that there could be room for more consistent and greater effects if this could be therapeutically driven.

An important discovery from this work was that the spinal neuron progenitor enriched donor cells give rise to additional cell populations after transplantation. New populations of spinal neurons, including inhibitory neurons and cells with more dorsally-associated genes, were detected 2 months post-transplantation. A crucial characteristic is that they were driven toward spinal fates prior to transplantation and retained that fate months later. They also gave rise to populations of glia for which there was no evidence from single-cell RNA-sequencing prior to transplantation. These glial and new neuronal populations, absent prior to transplantation, likely derived from the progenitor cells present in donor cells at the time of transplantation (clusters 1 and 3). The differentiation of transplanted progenitor cells is likely directed by the internal milieu of the injured spinal cord and by cytokines released by surrounding donor cells. The presence of both astrocytes and some oligodendrocytes is likely beneficial for the support of transplanted neurons, their integration with surrounding tissues and newly formed networks, and myelination.

Leveraging the wealth of advanced methods available to stem cell technologies, the field of neural transplantation is advancing at a rapid rate to provide more effectively tailored therapies for people living with injury and disease. Transplanting engineered V2a SpINs—a cell type known to contribute to plasticity and functional recovery—has offered a unique opportunity to use regenerative medicine to harness innate neural plasticity to enhance neural repair. As our knowledge of the cell types involved in neuroplasticity and repair continues to expand, so will the number of donor cell candidates that can be engineered for cell transplantation to enhance repair and recovery. Pairing more tailored donor cell therapies with complementary treatments, such as neural stimulation and rehabilitation, will enhance therapeutic efficacy and lasting functional recovery.

## Methods

### Animals

Total of 87 adult, female Sprague Dawley rats were used for this study (220-250g, Envigo). All animal procedures were approved by the Institutional Animal Care and Use Committee at Drexel University and National Institutes of Health guidelines for laboratory animal care and safety were strictly followed. Animals were housed in groups of 2-3 and had free access to food and water throughout the study, and had access to small enrichment items in each cage. All animals were housed in a 12h/12h light/day cycle, with surgeries and testing done during light cycles.

### Cell differentiation

Human induced pluripotent stem cells were grown to 70% confluency and passaged every 4 days as single cells onto Matrigel-coated cultureware. To seed cells for differentiation, single cells were plated at a density of 13,000 cells per cm^2^ with 10uM ROCK inhibitor, 10uM SB 431542, 0.1uL LDN193189 with varying concentrations of CHIR-99021 (0, 2, 4, 6uM) in mTeSR with a media change at day 3. At day 5, cells were fed with 100nM RA with same concentrations of LDN193189, SB 431542 and CHIR-99021 in Neural Induction Media (NIM). At day 7, cells were fed with NIM, RA, 1mM DAPT and 1mM Pur in NIM. Media was changed at days 10 and 13. At day 15, cells were lifted with accutase (15-minute incubations, three times with gentle trituration in between incubation periods), quenched with phosphate buffered saline and passed through 50um cell strainer prior to centrifugation (800rmp for 3 minutes), and resuspension in freezing medium for cryopreservation. Detailed methods and media components are available in Supplementary Methods.

### Real time quantitative polymerase chain reaction (RT-qPCR)

Samples were lysed within the 24-well plate and RNA was extracted using the E.Z.N.A. Total RNA Kit (Omega Biotek, Norcross, GA). RNA (500 ng) was reverse-transcribed into cDNA using the iScript cDNA synthesis kit (BioRad, Hercules, CA). RT-qPCR was performed using Fast SYBR Green Master Mix (Life Technologies) and the primers listed in Extended Data Table 5 were annealed at 61degC on the Step One Plus Real-Time PCR System (Life Technologies). Fold-changes were calculated using the ΔΔCt method.

### Immunocytochemistry

On Day 15 of differentiation, a subset (50,000 cells/coverslip) of cells lifted from 24-well plates were prepared for immunocytochemical analysis by plating onto poly-L-lysine and laminin coated 25mm^2^ glass coverslips and allowed to attach for 10 hours, 2 and 5 days (equivalent to Day 15, 17 and 20 in vitro) prior to fixation with paraformaldehyde (4% in PBS) for 10 minutes and staining. The cells were washed with PBS prior to blocking with normal donkey serum (GeneTex, #GTX73205) and 0.01% Triton (EMD Millipore, #9036-19-5) for one hour. Cells were stained with primary antibody (see Extended Data Table 6) overnight, washed three times with PBS, and stained with appropriate secondary antibody (see Extended Data Table 6). Cells were then washed three times for five minutes with PBS, prior to a 10-minute incubation with PBS containing Hoechst 33342 solution (1:10,000; ThermoFischer Scientific). Coverslips were secured onto slides with fluorescence mounting medium (Dako Agilent, #S302380-2) and imaged using a Zeiss Inverted Microscope attached to a Dell PC. Counting of Chx10 positive cells was performed in ImageJ software (NIH, Bethesda, MD) with approximately 200 cells counted (as determined by DAPI staining) per replicate (n=6).

### Multielectrode array

On Day 15 of differentiation, a subset (50,000 cells/well) were plated into poly(ethyleneimine) and laminin pre-coated wells of a 24-well CytoView multielectrode plate (Axion Biosystems, Atlanta, GA). Each well contains 16 PEDOT electrodes with a 350 μm electrode pitch. Plates were maintained in a 37°C/5% CO_2_ incubator and cells were fed every four days with neural maturation medium (BrainPhys plus SM1 supplement (Stemcell Technologies) supplemented with 10ng/ml of BDNF, GDNF, CNTF, and IGF, R&D Systems). Recordings were performed 1 day after feeding, using a Maestro Edge MEA system and Axis Navigator 2.0.4 software (Axion Biosystems). Data was acquired at 12.5 kHz, and spikes were detected in the AxIS Navigator software with an adaptive threshold crossing set to 6 times the standard deviation of the estimated noise for each electrode. Baseline data was acquired for 10 min at 37°C/5% CO_2_, prior to blue light stimulation at various frequencies (1, 1.5, 3Hz; Figure 1O). The last four minutes of each baseline recording were used for analysis, assessing bursting and network activity. Bursts were defined as events on individual electrodes with at least 5 spikes occurring with a minimum inter-spike interval of 100 milliseconds. Network bursts were defined as a minimum of 50 spikes with a 100 ms inter-spike interval and a minimum of 35% of the electrodes being considered active. Brightfield images of the cells or aggregates (Supplementary Figure 1) on the MEA plate were taken to assess electrode coverage.

### Spinal cord injury

All animals were prepared for aseptic surgery, underwent a laminectomy to expose the C3-4 spinal cord, and received a 200kD contusion injury using the Infinite Horizon Impactor. All animals went into respiratory arrest immediately or within 30 seconds of injury, and accordingly were mechanically ventilated for a maximum of 1 hour, at which point they were weaned from ventilation or excluded from the study if they did not recover voluntary control of breathing. A total of n=18 animals did not survive to the transplantation surgery 14 days post-injury.

### Cell preparation for transplantation

Cells were thawed from cryovials and suspended into 4mL Hanks Balanced Salt Solution, centrifuged at 800rmp for 3 minutes, and resuspended in 1mL of Hanks Balanced Salt Solution for cell counts. To prepare cells for injection, cells were centrifuged again and resuspended in Hanks Balanced Salt Solution to deliver desired dose (see below) of cells per 10uL. Cells were kept on ice until loading into a gas tight 25uL Hamilton syringe with a custom, 30-gauge needle, attached to a micromanipulator (Narishige) just before injection.

### Cell transplantation

Four days post-contusion, surviving animals began immunosuppression with Cyclosporin A, which they received daily for the duration of the study (10mg/kg, daily s.q.). Animals were blindly separated into vehicle treated and transplant recipients, and were decoded at the end of the study. One-week post-injury, animals were surgically anesthetized with a xylazine (10mg/kg s.q.) and ketamine (120mg/kg i.p.), and the skin and underlying musculature were re-exposed. Using a 33-gauge needle tip, a small hole was made in the dura overlying the center of the re-exposure and epicenter of the injury. Animals received either vehicle control (Hanks Balanced Salt Solution, HBSS) or V2a SpINs suspended in 8-10uL HBSS and pre-loaded into a gas tight 25uL Hamilton syringe. For the initial dosing study, animals received injections of either 250,000 (n=4), 500,000 (n=4) or 1 million cells (n=4) in 8-10 μL of HBSS, while remaining animals (n=58) received the highest dose of 1 million cells. Vehicle controls (n=17) received 8-10uL of HBSS into the lesion cavity. The underlying muscle was sutured in layers and the skin was closed with wound clips. Upon completion of each procedure anesthesia was reversed via injection of antisedan (1.2mg/kg s.q.). Post-operative medicine included lactated ringers (5mL, s.q.) to prevent dehydration and buprenorphine (0.025mg/kg s.q.) was used as an analgesic. Animals recovered for one month (cell dose study) or two-months prior to terminal electrophysiology recordings and perfusion.

### Transneuronal tracing

To anatomically trace the donor neurons that become integrated with the injured phrenic network, 50 μL of pseudorabies virus (PRV614 or PRV-Bartha, 2×10^8^ pfu) was topically applied to the hemi-diaphragm ipsilateral to injury, as described previously^26^.

### Brainstem AAV injections

One week prior to terminal electrophysiology, animals were secured in a stereotaxic frame and the caudal medulla was surgically exposed. Two 0.5 μL injections (World Precision Instruments syringe with 36 needle) of AAV1-Syn-Chrimson-tdT^21^ were made into the caudal medulla 1mm lateral to obex, and 1 and 1.5mm from the dorsal surface. The muscles were then sutured in layers and animals left to recover for one week to allow for anterograde transport of the td-Tomato and opsin through the axons of labeled bulbospinal neurons.

### Electrophysiology – diaphragm electromyography

Terminal electrophysiology was conducted in all animals, 2 months after transplantation. Animals were surgically anesthetized with ketamine and xylazine (as described above) for these recordings. All animals received a laparotomy and six electrodes were placed into three sites in the diaphragm for bipolar recordings of activity from the i) ventral diaphragm ipsilateral to injury, ii) dorsal ipsilateral diaphragm and iii) contralateral diaphragm. Spontaneous activity was recorded with the animal laying supine, under baseline conditions (breathing room air), and challenged conditions: exposed to five minutes of either hypoxic (10% O_2_ balanced in nitrogen) or hypercapnic gas (7% carbon dioxide, 20% oxygen, balanced in nitrogen).

### Electrophysiology – intraspinal stimulation and multiunit recording

A subset of animals received blue or red-light stimulation and multi-unit recordings from the transplant. For these recordings, the animals’ abdomen was stapled closed and the animal was turned to lay in the prone position. The site of the transplanted was re-exposed and a fiber optic cannula (0.5mm, Thor labs) was placed on the surface of the spinal cord at the site of injury and transplant. A carbostar-1 electrode (Kation scientific) was then inserted 0.5 mm into the site of the transplant immediately adjacent to the fiber optic. Spontaneous donor neuron (multi-unit) activity was recorded and a new baseline EMG recording was obtained from this prone position. Once baseline was completed, blue or red light was delivered through the cannula using a High Power LED Driver (Thor labs DC2100, 1000mA current, 50% duty cycle, at 1, 3, 20 and 50Hz) to activate channelrhodopsin-expressing donor cells or any AAV-Chrimson labeled axons innervating the transplant, respectively. Given the limited tissue penetration of blue light, if there were no observable changes in multi-unit activity from the transplant during blue light stimulation from the top of the spinal cord, the fiber optic cannula was placed into the spinal tissue 1mm from the dorsal surface and the stimulation was performed again.

### Tissue collection

Upon completion of electrophysiological studies, the animals were euthanized and either intracardially perfuse-fixed with paraformaldehyde (4% w/v in 0.1M PBS), or animals prepared for fresh tissue removal and subsequent isolation of nuclei, as described in detail previously^27^.

### Immunohistochemistry

Brains and spinal cords of animals that were fixed via intracardial perfusion were post-fixed in paraformaldehyde (4% w/v in 0.1M phosphate buffered saline (PBS), pH 7.4) for 1-3days. Spinal cord tissue spanning caudal-most part of brainstem to beginning of thoracic spinal cord (T1-2) or brainstem were cryoprotected (15%, then 30% sucrose, overnight), and sectioned (on-slide 30µm or free floating 40µm, longitudinally) using a cryostat. On slide sections were rehydrated for 15 minutes in PBS, blocked against endogenous peroxidase activity (30% methanol, 0.6% hydrogen peroxide in 0.1M PBS, incubated for 1 hour, 2 hours for free floating sections), and blocked against non-specific protein labeling (10% serum in 0.1M PBS with 0.02% Triton-X, incubated for 1 hour, 2 hours for free floating sections), prior to application of primary antibodies in blocking solution. Primary antibodies (summarized in Extended Data Table 6) were left on the tissue overnight at room temperature. Antibodies to PRV were supplied by Dr. Lyn Enquist (Princeton University). The following day, tissue was washed in PBS (0.1M, 3 x 5 minutes) and incubated in blocking solution with secondary antibodies (summarized in Extended Data Table 6). Immunolabeled sections were then washed in PBS (0.1M, 3 x 5 minutes), allowed to dry, and coverslipped with fluorescence mounting medium (Dako). Sections were examined using a Zeiss Inverted microscope, attached to a Dell PC. Photographs were taken with a digital camera (Zeiss Axiocam MRm).

### Single cell and nuclear RNA sequencing

For single cell sequencing, cells previously cryopreserved at day 15 of culture were rapidly thawed and approximately 40,000 cells were prepared for single cell analysis. For single nuclear sequencing, animals were euthanized and nuclei were isolated from fresh spinal cord tissue as previously described^27^. Cells and nuclei were prepared by droplet encapsulation with the Chromium Controller and library preparation with the Chromium Single Cell 3’ v1 Library and Gel Bead Kit according to the manufacturer’s instructions (10x Genomics, San Francisco, CA). Final libraries were sequenced on a NextSeq500 (Illumina, San Diego, CA).

### Custom reference genome

A custom mouse reference genome was created using the reference human genome sequence (GRCh38) from Ensembl (release 98)^28^ and the human gene annotation file from GENCODE (release 32)^29^, similar to those used in 10x Genomics Cell Ranger human reference package GRCh38 2020-A. The headers of the Ensembl reference mouse genome sequence fasta file with the chromosome names were modified to match the chromosome names in a fasta file from GENCODE. The annotation GTF file contains entries from non-polyA transcripts that overlap with the protein coding genes. These reads are flagged as multi-mapped and are not counted by the 10x Genomics Cell Ranger v.6.1.1 count pipeline^30^. To avoid this, the GTF file was modified to (1) remove version suffixes from transcript, gene and exon IDs to match the Cell Ranger reference packages, and (2) remove non-polyA transcripts. The sequence for marker genes (YFP and ChR2) were appended as separate chromosomes to the end of the human reference genome sequence and the corresponding gene annotations were appended to the filtered human reference gene annotation GTF file. The 10x Genomics Cell Ranger v.6.1.1 mkref pipeline was used to build the custom reference genome using the modified fasta and GTF file.

### Pre-processing and clustering of single-cell RNA-seq data

The scRNA-seq data included a single sample split into 4 lanes for sequencing. The demultiplexed fastq files for these lanes were aligned to the custom human reference genome (custom reference genome methods provides additional descriptions) using the 10x Genomics Cell Ranger v.6.1.1 count pipeline^30^, as described in the Cell Ranger documentation. The filtered count matrices generated by the Cell Ranger count pipeline were processed using the R package Seurat v.4.1.1^31^. Each lane was pre-processed as a Seurat object such that only cells with at least 200 genes detected and features detected in at least 3 cells were kept. The top 1% of cells per sample with a high number of unique genes, cells with ≤750 unique genes and cells ≥5% mitochondrial genes were filtered out. The pre-processed data for 4 lanes were merged into a single Seurat object and normalization and variance stabilization was performed using sctransform^32^ with the ‘glmGamPoi’ (Bioconductor package v.1.6.0) method^33^ for initial parameter estimation. Graph-based clustering was performed using the Seurat v.4.1.1 functions Find Neighbors and Find Clusters. First, the cells were embedded in a k-nearest neighbor graph based on the Euclidean distance in the principal-component analysis space. The edge weights between two cells were further modified using Jaccard similarity. Clustering was performed using the Louvain algorithm implementation in the FindClusters Seurat function. Clustering was performed for all combinations of 10, 15 and 20 principal components (PCs) with 0.01,0.02,0.03,0.04,..,0.2,0.1, 0.2, 0.3, 0.4, 0.5 and 0.6 resolutions. Clustering with 20 PCs and 0.09 resolution resulted in 5 distinct biologically relevant clusters.

### Cell type assignment

Data visualization using Seurat v.4.1.1 in the UMAP space for the 4 lanes revealed no batch effects by sequencing date. The marker genes for each cluster were identified using the FindAllMarkers Seurat function on the SCT assay data. This algorithm uses the Wilcoxon rank-sum test to iteratively identify differentially expressed genes in a cluster against all the other clusters. Marker genes were filtered to keep only positively expressed genes, detected in at least 25% of the cells in either population and with at least 0.25 log_2_ fold change. We assigned identities to cell clusters by matching the cell clusters to known cell types with the expression of canonical cell-type-specific genes.

### Pre-processing and clustering of single-nuclear RNA-seq data

The snRNA-seq data included a single sample split into 4 lanes for sequencing. The demultiplexed fastq files for these lanes were aligned to the custom human reference genome (custom reference genome methods provides additional descriptions) using the 10x Genomics Cell Ranger v.6.1.1 count pipeline^30^, as described in the Cell Ranger documentation. The filtered count matrices generated by the Cell Ranger count pipeline were processed using the R package Seurat v.4.1.1^31^. Each lane was pre-processed as a Seurat object such that only cells with at least 200 genes detected and features detected in at least 3 cells were kept. The top 1% of cells per sample with a high number of unique genes, cells with ≤1,000 unique genes and cells ≥0.2% mitochondrial genes were filtered out. The pre-processed data for the 4 lanes was merged into a single Seurat object and normalization and variance stabilization was performed using sctransform^32^ with the ‘glmGamPoi’ (Bioconductor package v.1.6.0) method^33^ for initial parameter estimation. Graph-based clustering was performed using the Seurat v.4.1.1 functions FindNeighbors and FindClusters. First, the cells were embedded in a k-nearest neighbor graph based on the Euclidean distance in the principal-component analysis space. The edge weights between two cells were further modified using Jaccard similarity. Clustering was performed using the Louvain algorithm implementation in the FindClusters Seurat function. Clustering was performed for all combinations of 10, 15 and 20 principal components (PCs) with 0.01,0.02,0.03,0.04,..,0.2,0.1, 0.2, 0.3, 0.4, 0.5 and 0.6 resolutions. Clustering with 20 PCs and 0.6 resolution resulted in 20 distinct biologically relevant clusters for the snRNA-seq data.

### Combined analysis of single-cell RNA-seq data and single-nuclear RNA-seq data

The pre-processed data for both scRNA-seq and snRNA-seq samples from all 4 lanes was merged into a single Seurat object. The Seurat object was processed in 3 different ways: (1) normalization and variance stabilization was performed using sctransform^32^ with the ‘glmGamPoi’ (Bioconductor package v.1.6.0) method^33^ for initial parameter estimation, (2) normalization and variance stabilization was performed using sctransform^32^ with the ‘glmGamPoi’ (Bioconductor package v.1.6.0) method^33^ for initial parameter estimation while regressing out the effect of cell cycle genes and the number of genes detected in each cell, (3) normalization and variance stabilization was performed using sctransform^32^ with the ‘glmGamPoi’ (Bioconductor package v.1.6.0) method^33^ for initial parameter estimation followed by harmony correction for the data type (i.e. single-cell vs single-nuclear RNA-seq data). Graph-based clustering was performed using the Seurat v.4.1.1 functions FindNeighbors and FindClusters. First, the cells were embedded in a k-nearest neighbor graph based on the Euclidean distance in the principal-component analysis space. The edge weights between two cells were further modified using Jaccard similarity. Clustering was performed using the Louvain algorithm implementation in the FindClusters Seurat function. Clustering was performed for 20 principal components (PCs) and 0.09 resolution. All 3 normalization and variance stabilization methods resulted in clustering by data type with a few cells from the two data types with similar expression profiles.

### Statistical analysis

Statistical analyses were performed using Prism 9 (GraphPad, San Diego, CA) software with designated significance level of 95%. Data are presented as mean ± s.e.m with individual data scatter plots. One-way ANOVA was used for comparisons with Tukey’s post-hoc when the F-test revealed the existence of significant difference between tested groups.

## Acknowledgements

We thank members of the Srivastava, Lane and McDevitt laboratories for discussion and feedback of these studies. We thank the Gladstone Editing Team, Francoise Chanut and Kathryn Claiborn, for their feedback and revisions of the manuscript. We acknowledge the UCSF Center for Advanced Technology (CAT, supported by UCSF PBBR, RRP IMIA, and NIH 1S10OD028511-01 grants). We thank Mylinh Bernardi and Horng-Ru Lin of the Gladstone Genomics Core for their assistance with planning and execution of library preparation for single cell and nuclear sequencing. We thank the Gladstone Stem Cell Core and Gladstone Histology and Microscopy Cores for their technical support and the Drexel University Laboratory Animal Resources (ULAR) facility at Queen Lane College of Medicine for support with animal housing and care. We thank the NIH Virus Center grant no. P40 OD010996 and David Bloom at the University of Florida for pseudorabies virus studies.

LVZ was supported by the Lisa Dean Moseley Foundation, NIH NINDS F32 NS119348 and CIRM DISC2-14180. TF was supported by NIH NINDS F31 NS125975. MW was supported by the STAR Scholars Program at Drexel University. TM was supported by the Roddenberry Foundation. MAL was supported by R01 NS104291 and the Lisa Dean Moseley Foundation. DS was supported by the Roddenberry Foundation. OFV was supported by UCSF Health Innovation via Engineering (HIVE) initiative.

## Author Contributions

LVZ, MAL, TM, DS conceived the study, interpreted the data and wrote the manuscript. LVZ, MAL, TF, MW performed survival and terminal surgeries and animal care. LVZ and OFV performed cell differentiations and multielectrode recordings. LVZ performed all molecular qPCR, immunocytochemistry, immunohistochemistry, microscopy and analysis. LVZ performed all electrophysiological data analysis. AA performed single-cell and single-nuclear RNA-sequencing analysis.

## Additional information

Correspondence and requests for materials should be addressed to L.V.Z. or D.S. Reprints and permissions information is available at http://www.nature.com/reprints. Extended Data Information is available for this paper.

---

**Extended Data Figure 1.**
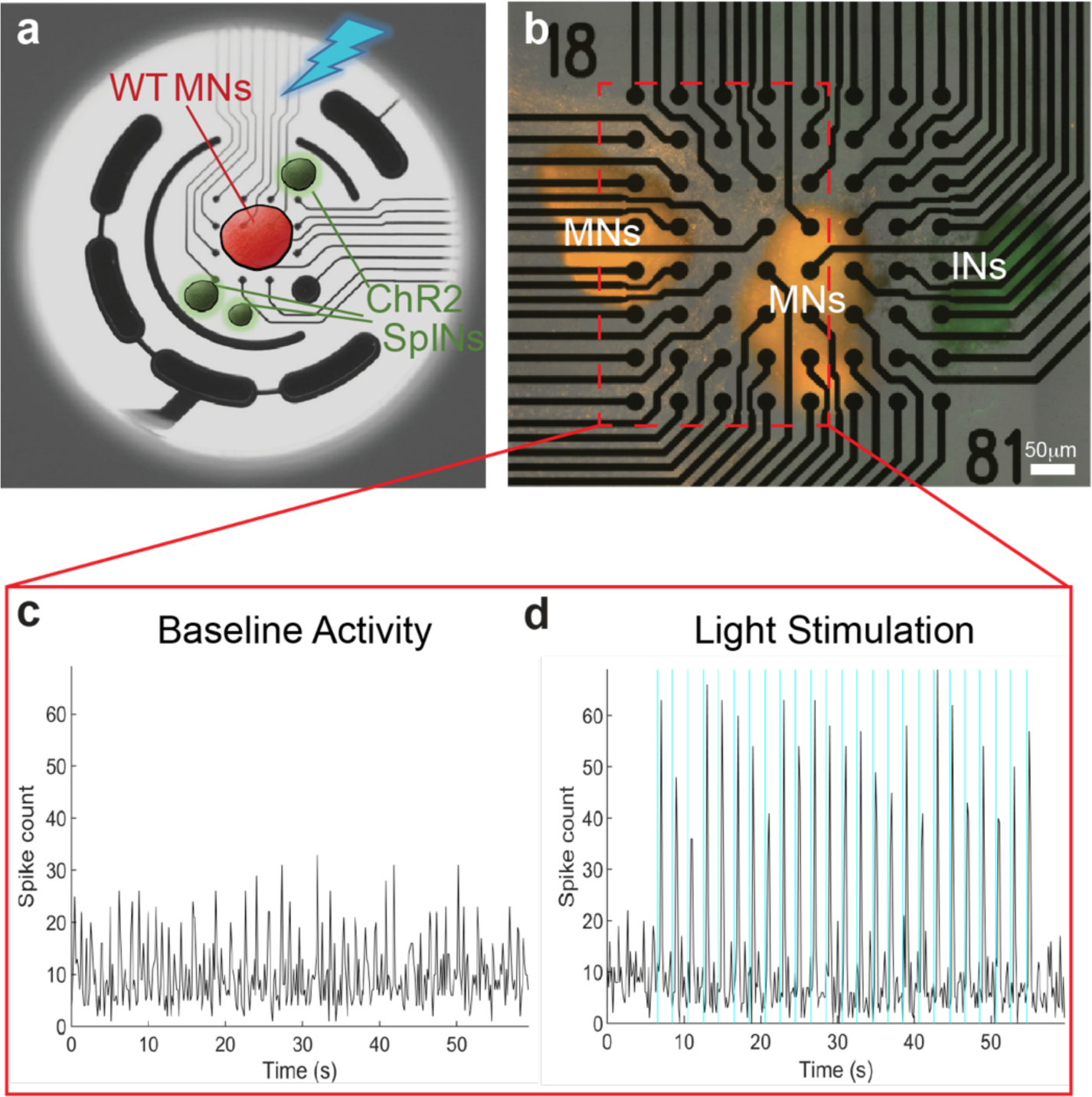
Human SpINs that were engineered to express the channel rhodopsin receptor and green fluorescent protein, were plated on multielectrode arrays (MEAs) with nearby aggregates of spinal motor neurons (MNs) expressing red fluorescent protein, as shown in the schematic diagram (a) and brightfield image (b). Blue light stimulation of optogenetic SpINs resulted in measurable activation of MNs (d compared to baseline activity in c), demonstrating formation of functional synapses between the SpINs and MNs. MNs: motor neurons; SpINs: spinal interneurons.

**Extended Data Figure 2.**
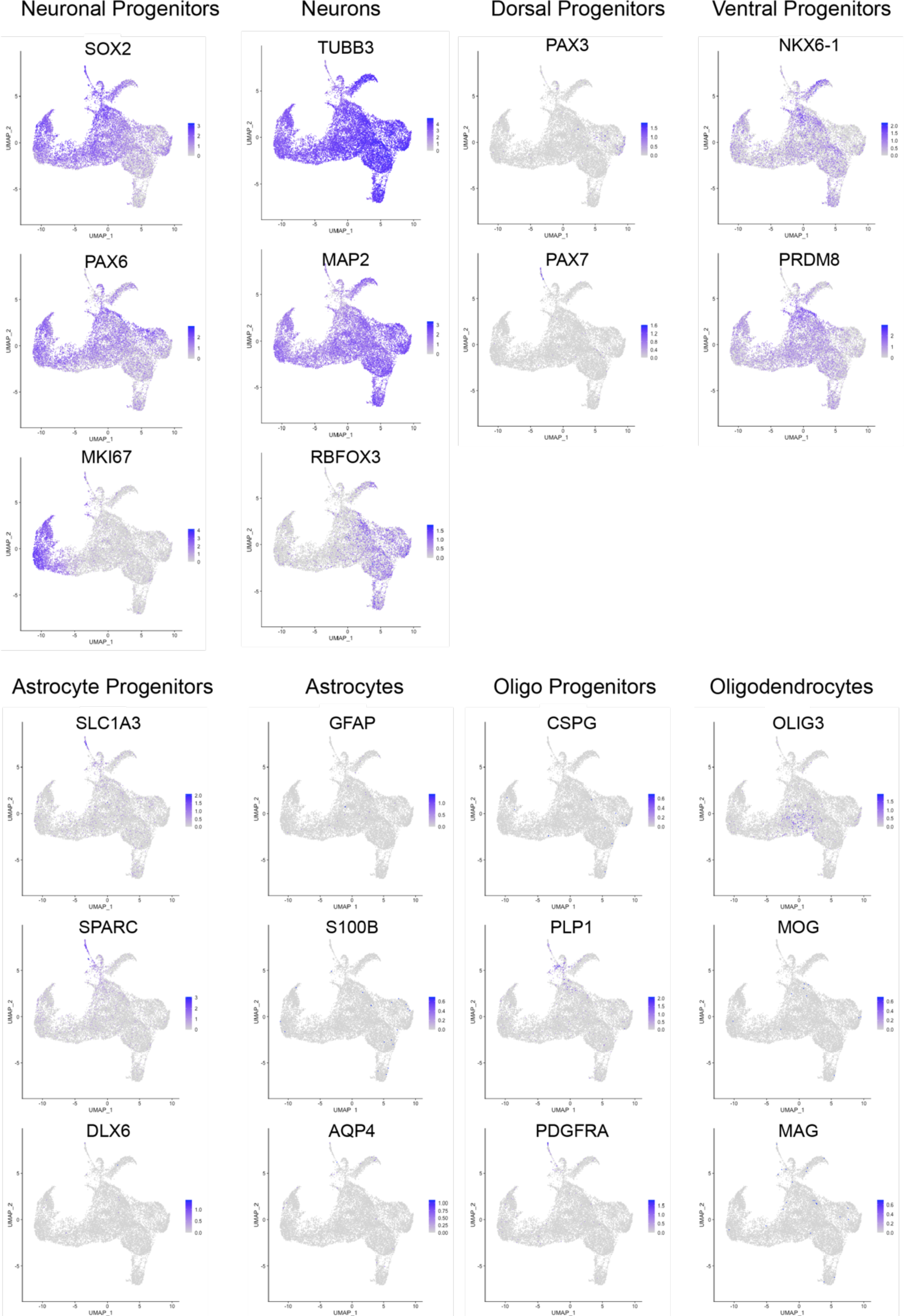
UMAPs of neuronal and glial markers expressed in Day 15 hiPSCs differentiated toward spinal interneurons analyzed with single cell sequencing. Both mature and progenitor neuronal markers were identified, with the latter being predominantly from ventral progenitors. Few astrocyte and oligodendrocyte markers were identified.

**Extended Data Figure 3.**
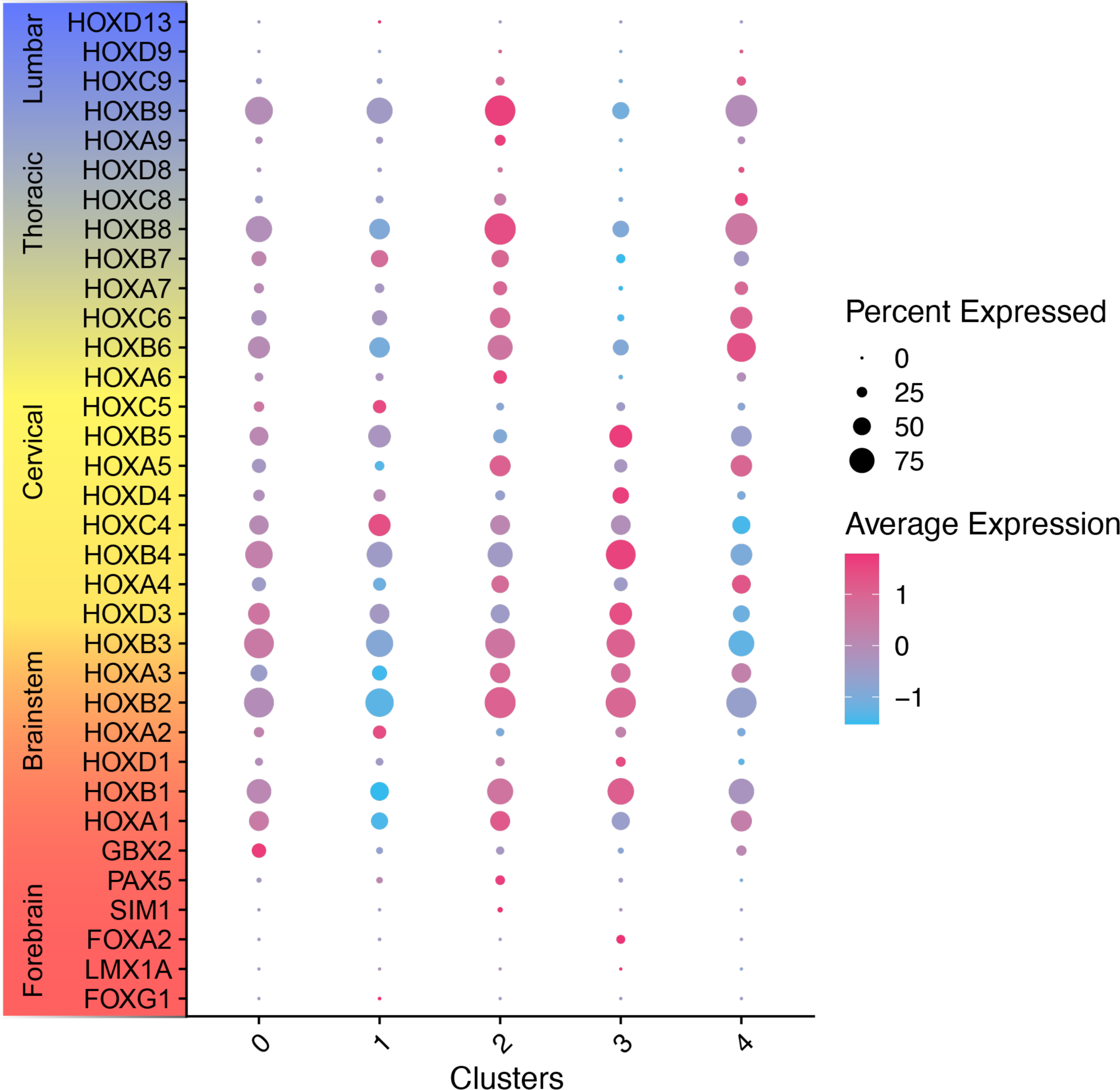
Hox Gene Expression in hiPSC-derived SpINs. Dot plot representing HOX gene expression for each of the clusters identified from scRNAseq of Day 15 hiPSCs differentiated toward SpINs. Note that cells within the cervical spinal cord express HOX genes 3-8.

**Extended Data Figure 4.**
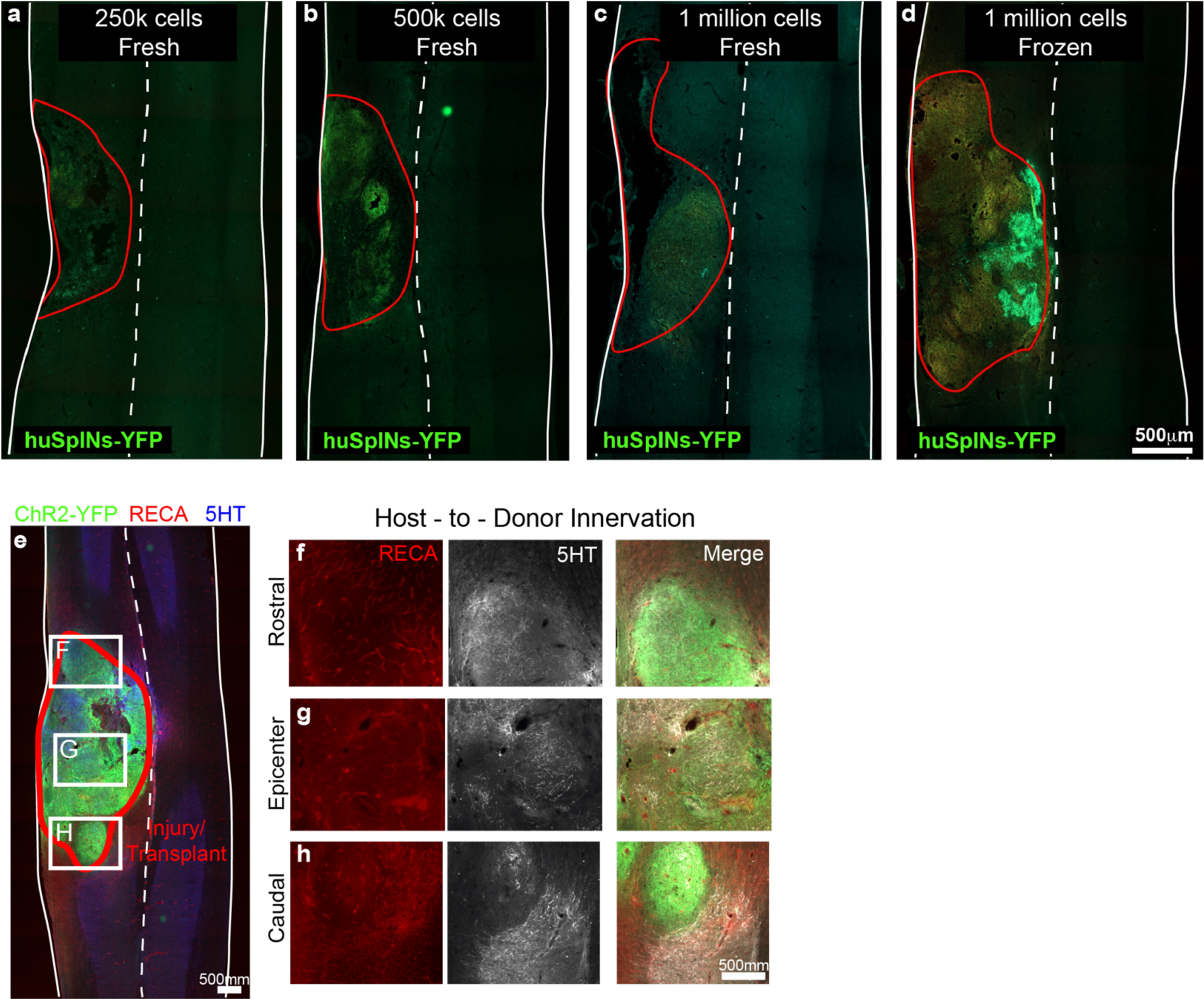
Coronal sections of the cervical spinal cord show the site of spinal cord injury and transplanted cells, 1-month post-transplantation. Fluorescence of transplanted ChR2-YFP-labeled SpINs at doses indicated (a-d) was performed to assess the cell dose that resulted in best donor cell survival, and greatest host-donor-host confluency (d). Fresh cells dosed at 250k, 500K and 1 million cells (thawed and cultured for 3 days prior to transplantation) and 1 million frozen cells (thawed and transplanted immediately and considered ‘off-the-shelf’). Immunocytochemistry for rat endothelial cell antigen (RECA) and serotonin (5HT) 2 months after transplantation labeled host endothelial cells and serotonergic axons, respectively, integrated with transplanted cells (YFP) (e-h).

**Extended Data Figure 5.**
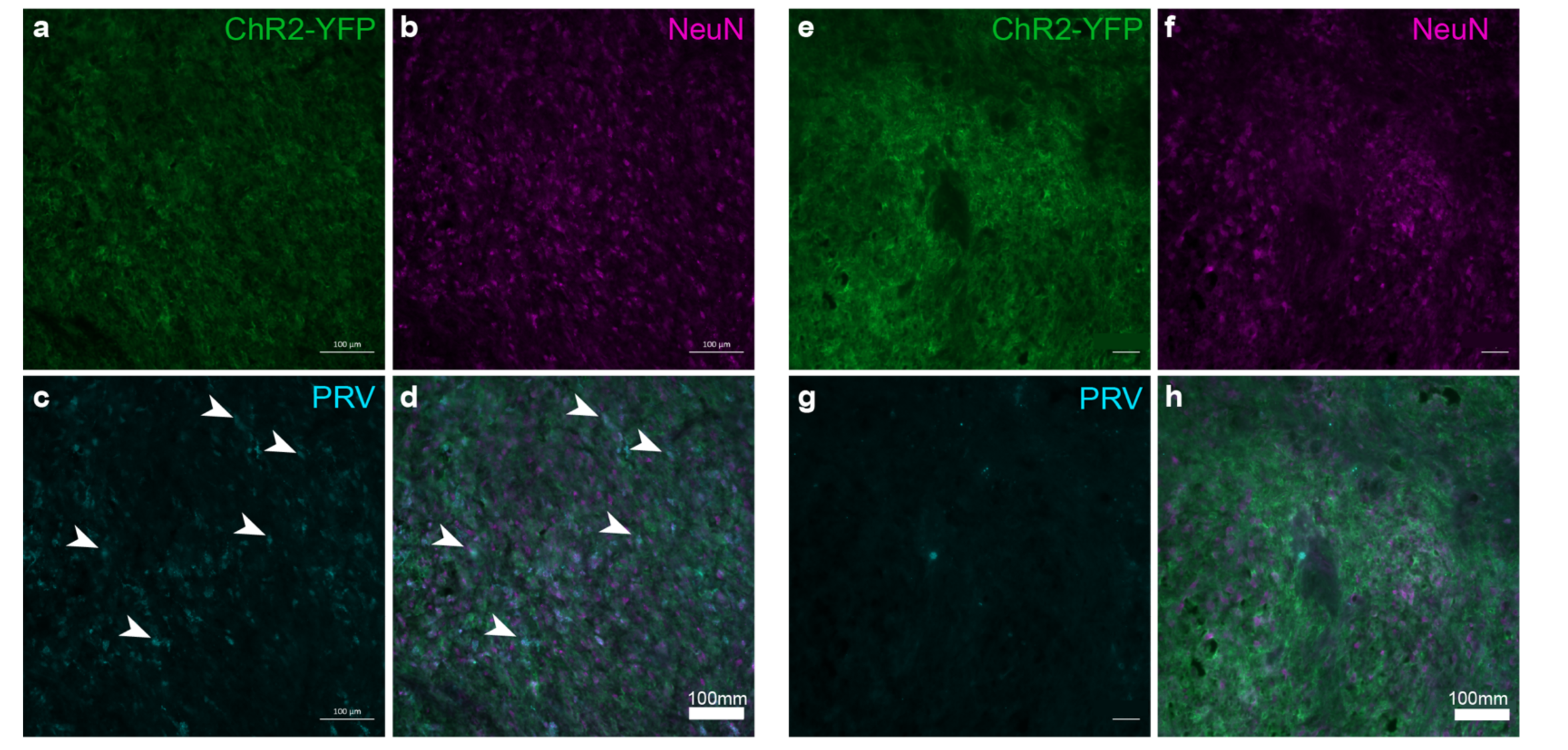
Coronal sections of transplant epicenter (ChR2-YFP, a, e) immunohistochemically labeled for neurons (NeuN, b, f) and pseudorabies virus (PRV, c, g) reveals distinct labeling patterns in two examples from independent rats (merge in d, h). White arrowheads point to PRV+ donor cells that are synaptically integrated with the phrenic network ipsilateral to injury.

**Extended Data Figure 6.**
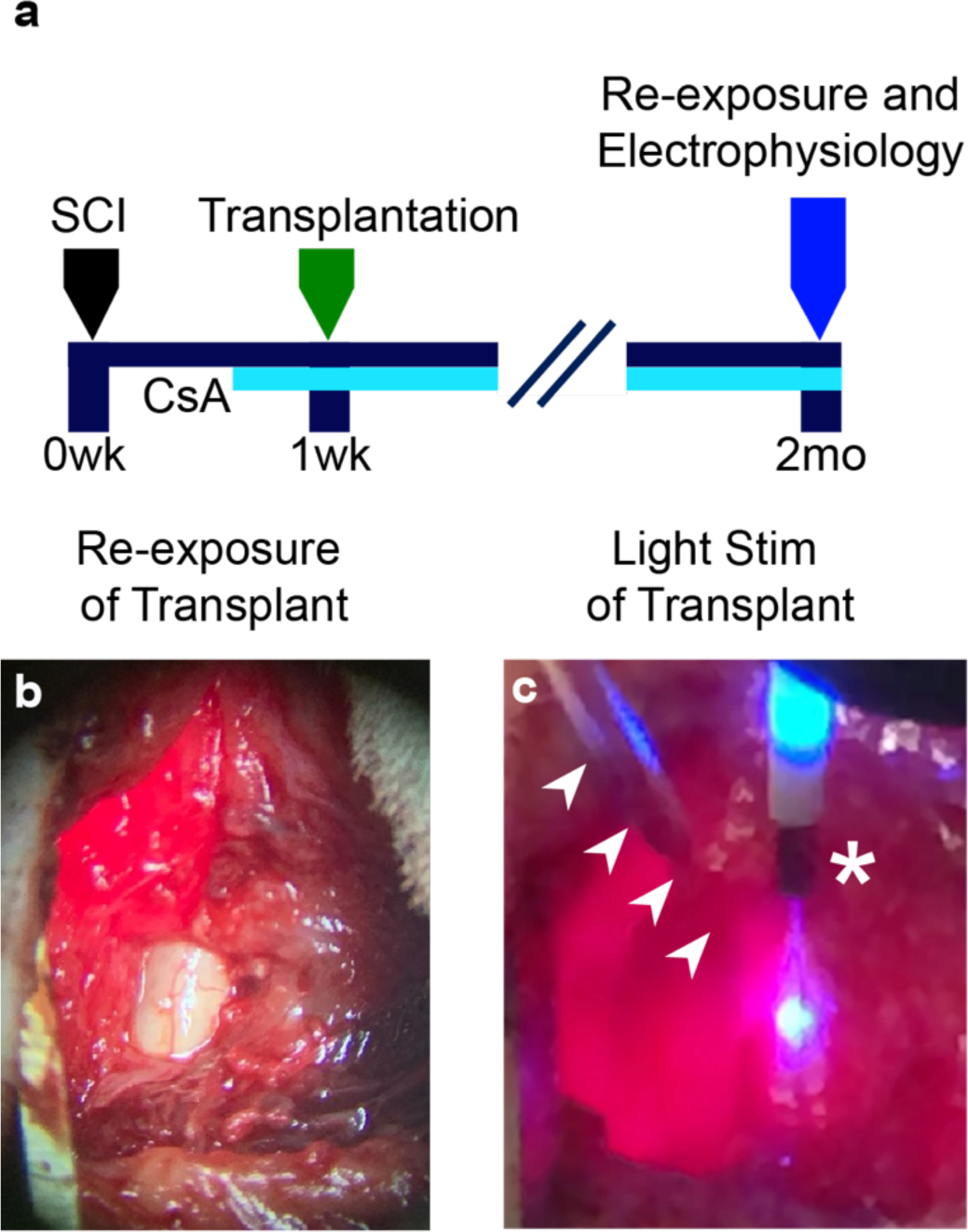
Experimental timeline (a) and images of re-exposed site of transplant (b) 2-months post-transplantation to enable recording and light stimulation of transplanted cells (c). White arrowheads indicate Carbon electrode (Kation Scientific) implanted into transplant. White asterisk indicates cannula (ThorLabs) delivering blue light to the transplant.

**Extended Data Figure 7.**
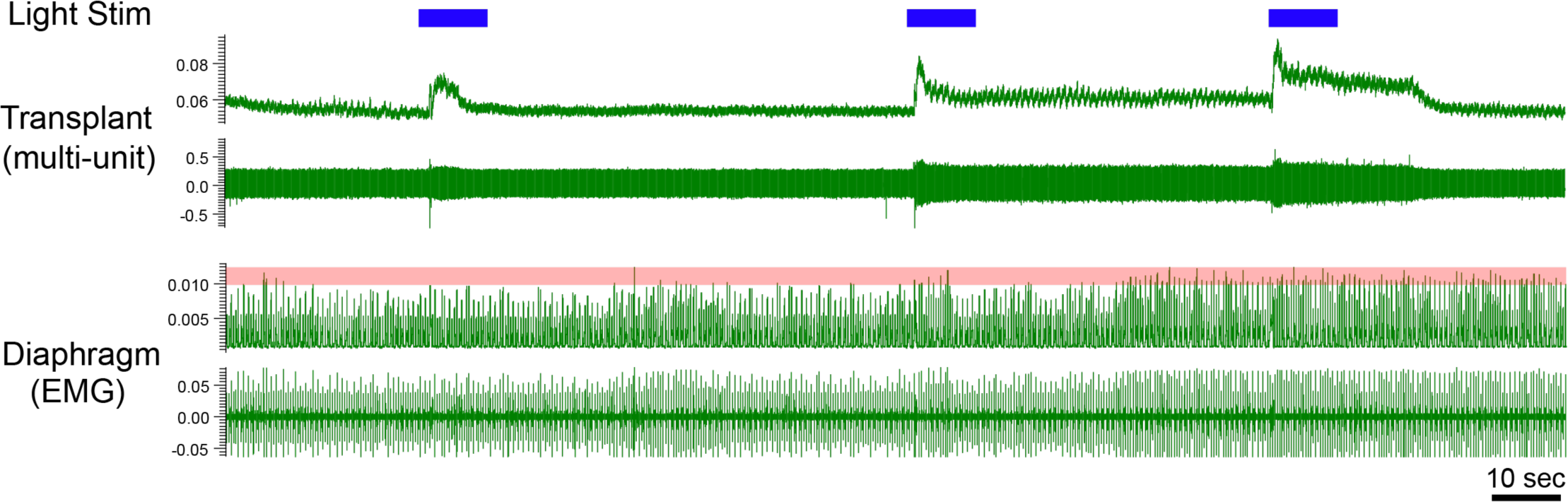
Examples of multi-unit and muscle myogram recordings from the transplant and diaphragm, respectively (raw and integrated traces are on bottom and top, respectively). Optogenetic stimulation of transplanted cells (blue bar) was repeated to assess whether the response to stimulation was reproducible.

**Extended Data Figure 8.**
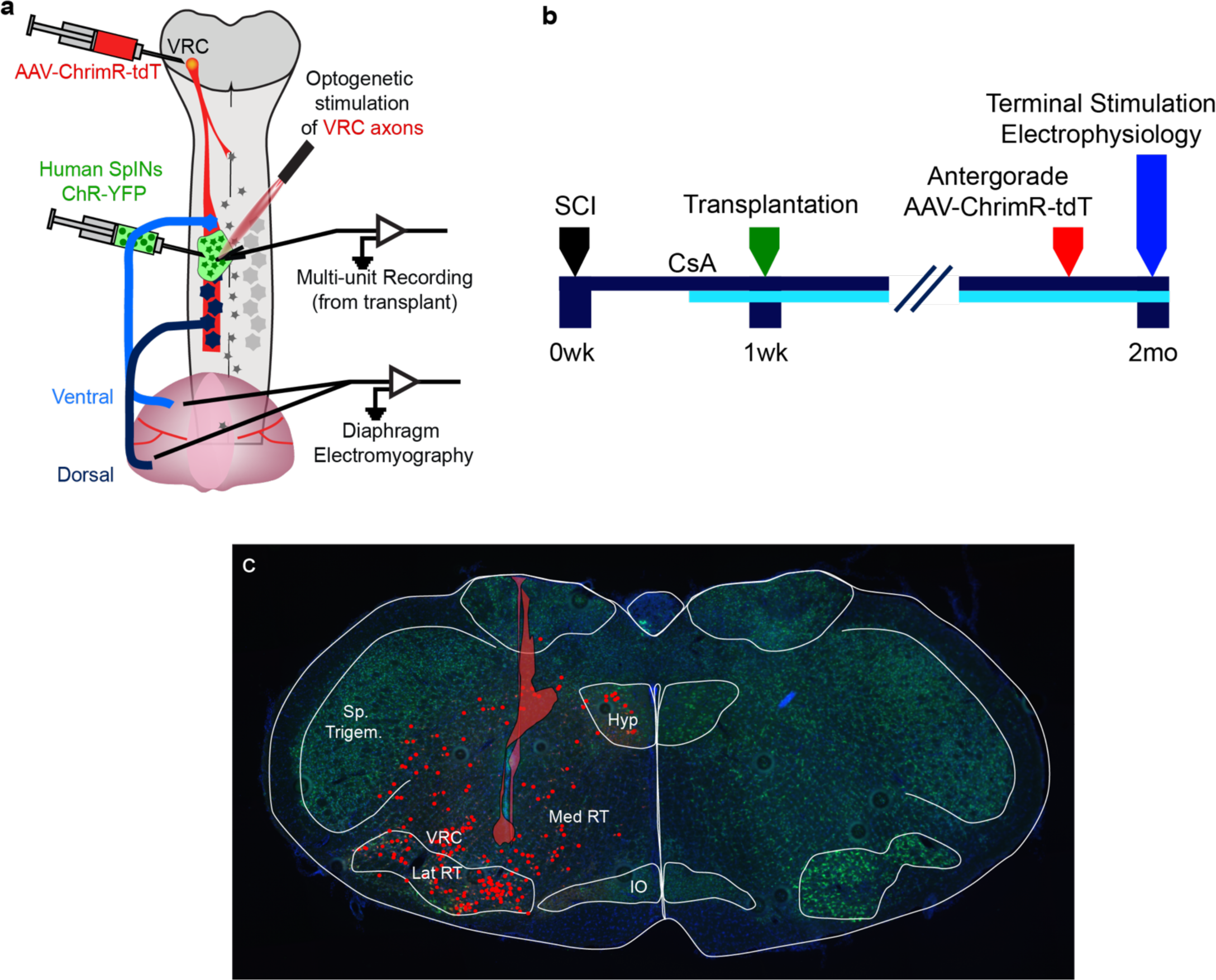
Schematic diagram of experimental design (a) and timeline (b) for delivery of AAV-ChrimR-tdT to brainstem respiratory centers. Two months post-transplantation and 1 week prior to terminal electrophysiology, injections were peformed into the reticular nuclei of the caudal medulla rostral to injury and transplant to assess the functional connectivity between host neurons and transplanted human spinal interneurons (Human SpINs ChR-YFP). A cross-section through the medulla (c; identifiable cytoarchitecture traced in white) at the site of injection shows the site of injection (red overlay trace, n=3 animals) and the distribution of AAV-labeled cells (red dots). The reticular nuclei (the ventral respiratory column (VRC), lateral and medial reticular nuclei (Lat RT, Med RT) have bulbospinal projections to spinal networks, and the VRC is responsible for providing respiratory drive to the phrenic network in the intact and injured spinal cord. Spinal trigeminal nucleus (Sp. Trigem.), hypoglossal nucleus (Hyp), inferior olive (IO), medial and lateral reticular nuclei (MedRT, LatRT), ventral respiratory column (VRC).

**Extended Data Figure 9.**
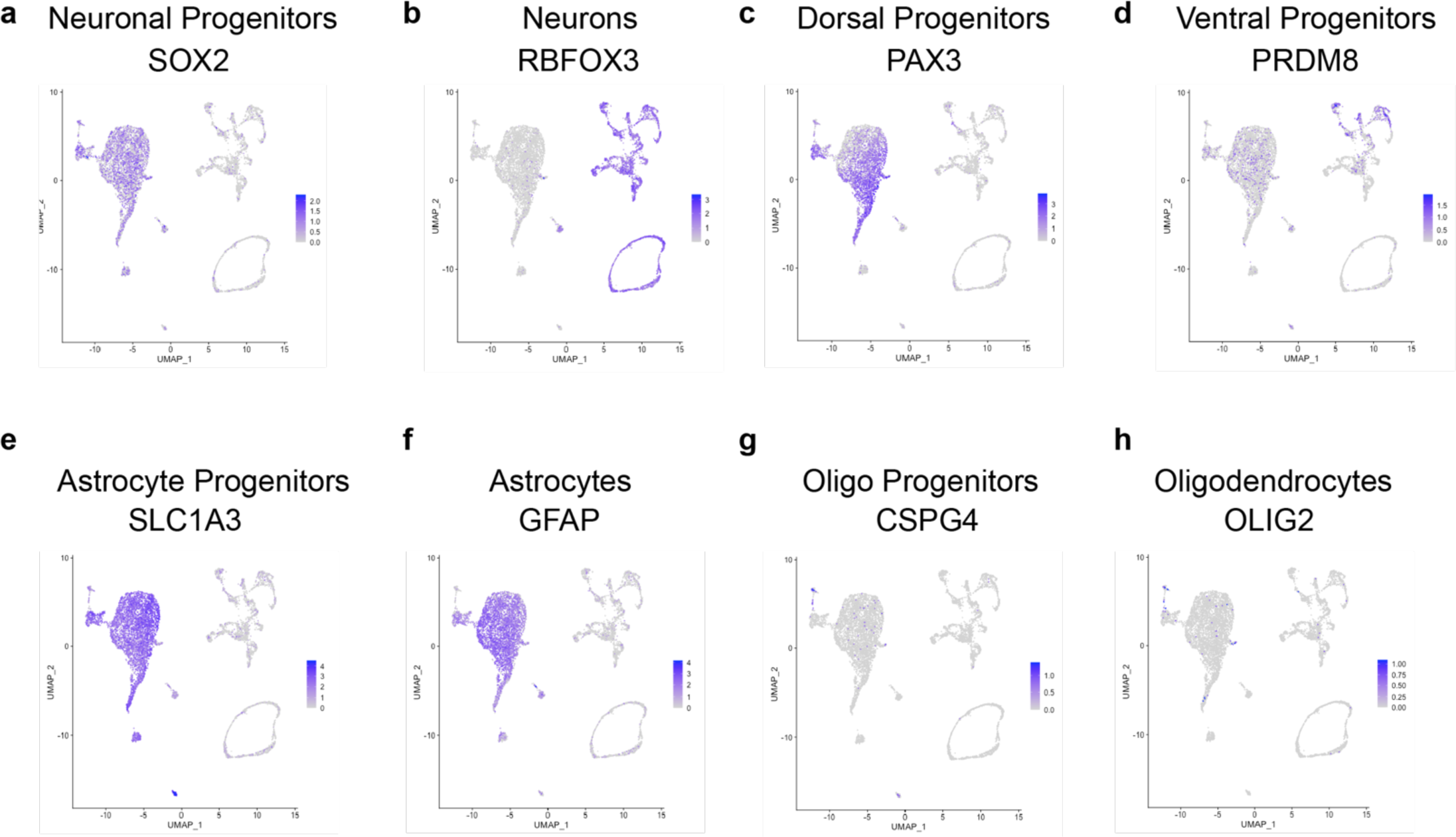
UMAPs of gene expression typical of neuronal progenitors (a), neurons (b), dorsal (c) and ventral (d) progenitors, as well as astrocyte progenitors (e), astrocytes (f), oligodendrocyte progenitors (g) and oligodendrocytes (h) in nuclei isolated from donor cells 2 months after being transplanted into the injured rat spinal cord. Compared to the single-cell sequencing data from donor cells prior to transplantation, there is an increase in glial markers (especially astrocytic) and dorsal neuron progenitors.

**Extended Data Figure 10.**
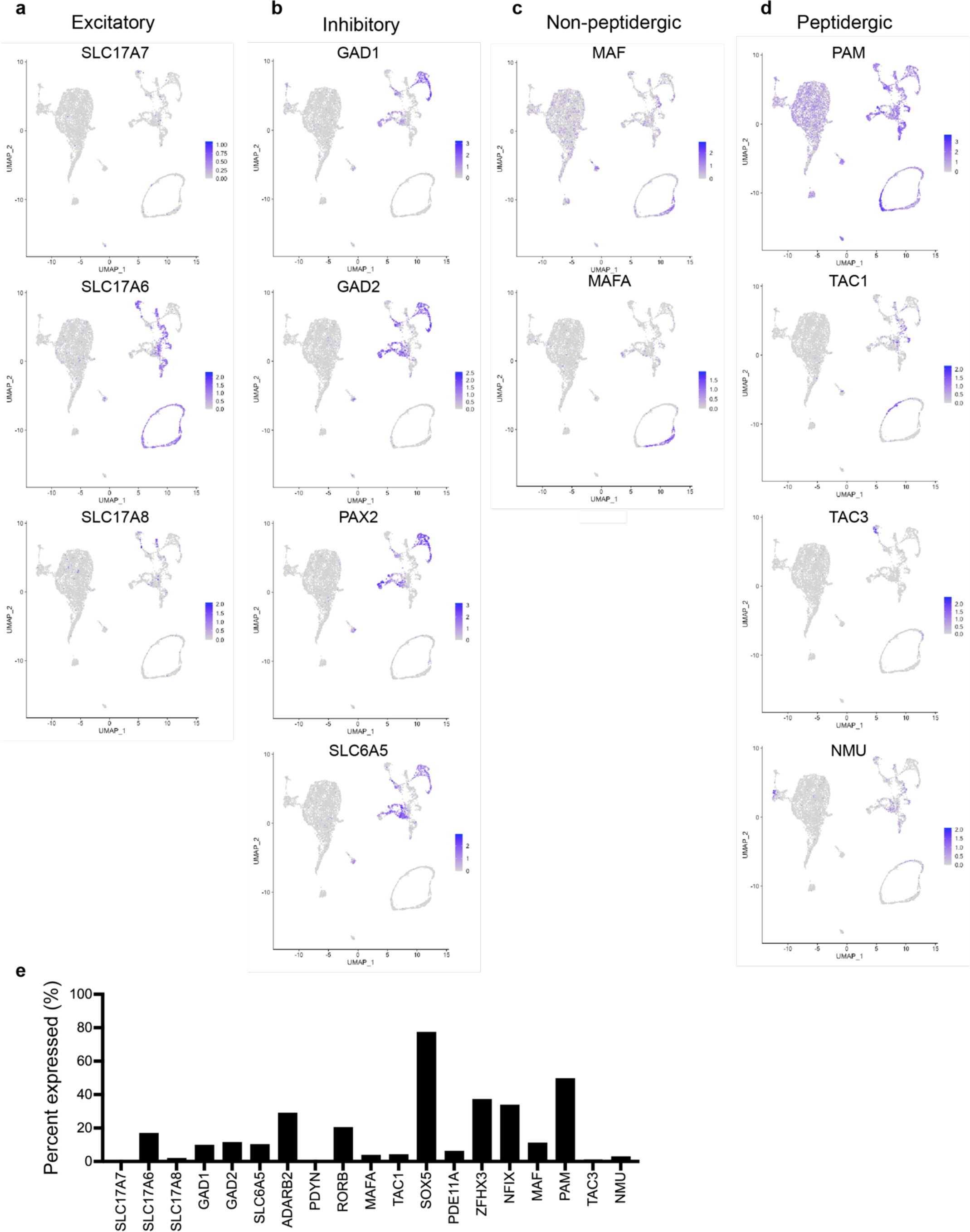
UMAPs of genes expressed in nuclei isolated from donor cells 2-months post-transplantation, showing presence of excitatory (a), inhibitory (b), non-peptidergic (c) and peptidergic (d) neuronal subtypes. Relative expression of each lister neuronal gene (e).

**Extended Data Figure 11.**
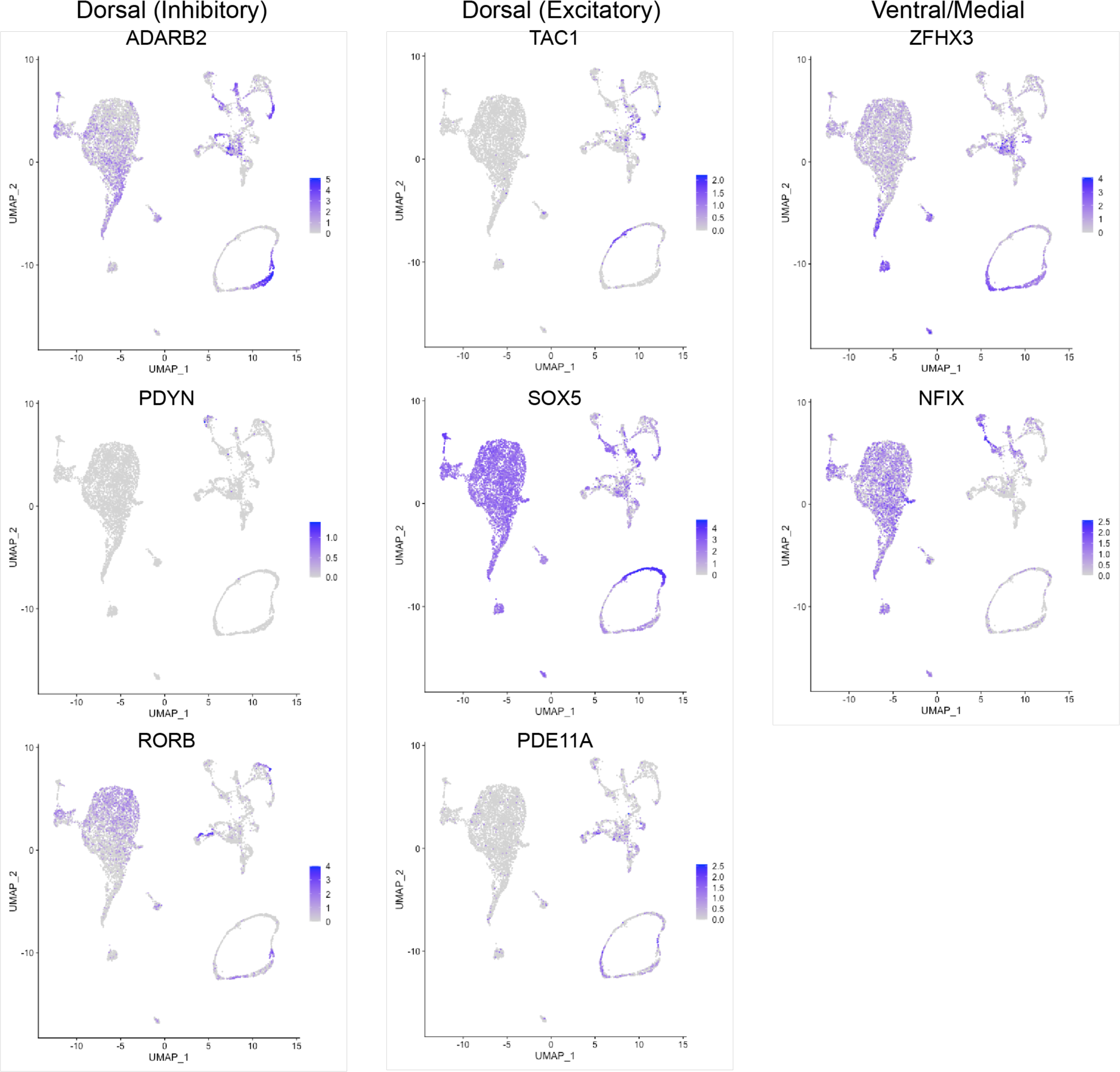
UMAPs of genes expressed in nuclei isolated from donor cells 2-months post-transplantation, showing expression of markers related to diverse population of neurons.

**Extended Data Figure 12.**
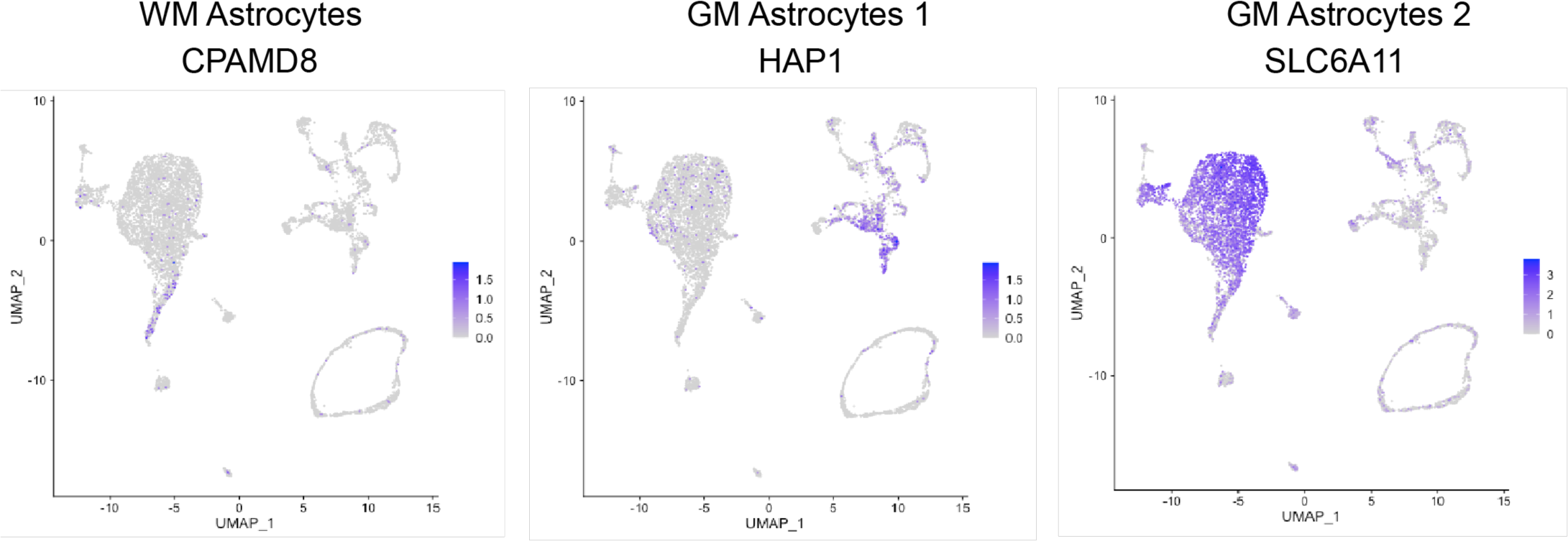
UMAPs of gene expression typical of white and gray matter astrocytes sequenced from nuclei isolated from donor cells 2 months after being transplanted into the injured rat spinal cord.

**Extended Data Figure 13.**
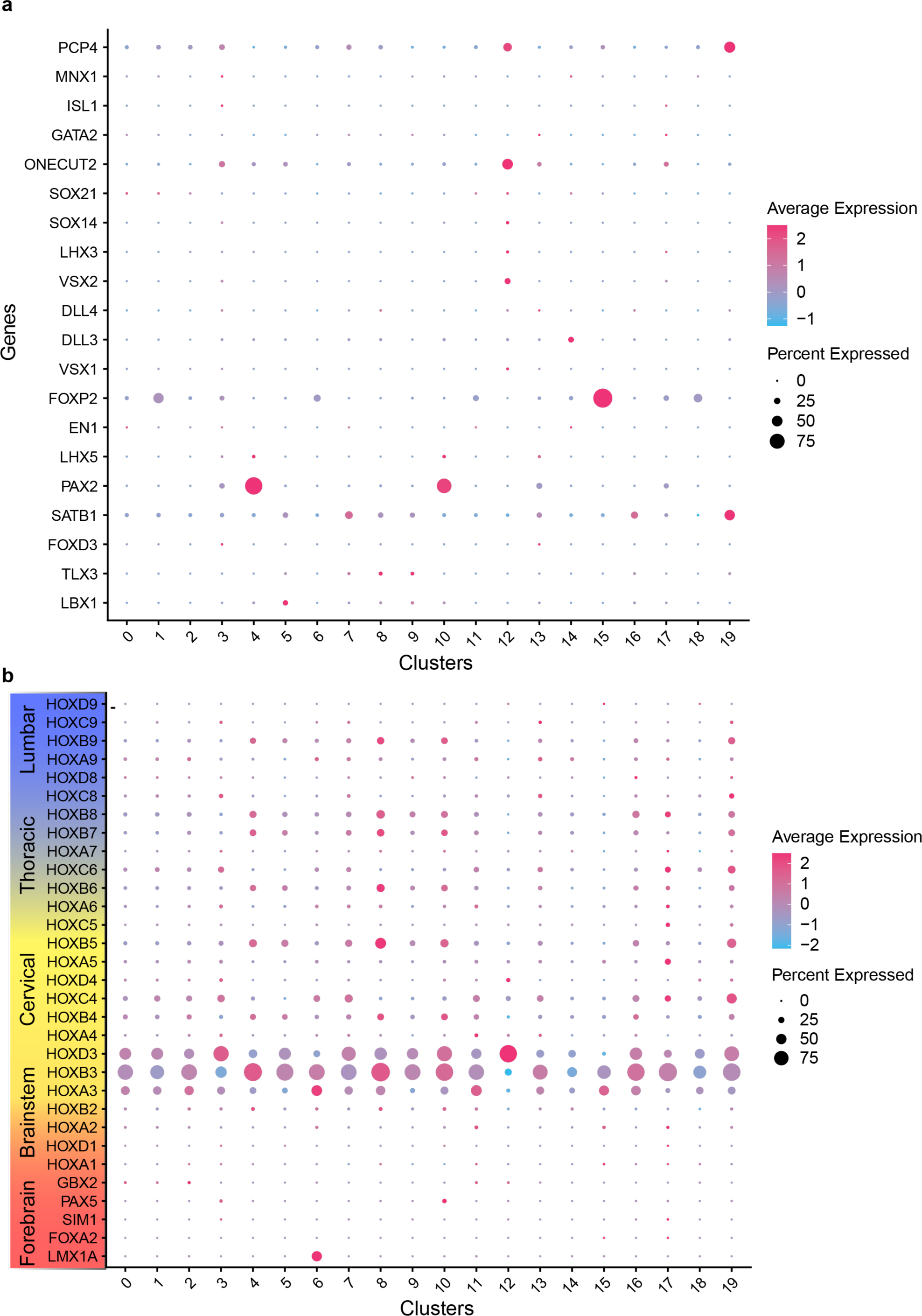
Dot plot of all hox genes (a) and developmental transcription factors (b) expressed in nuclei isolated from donor cells 2-months post-transplantation into injured spinal cord. Note that cells within the cervical spinal cord express HOX genes 3-8.

